# Tracing the evolution of the plastome and mitogenome in the Chloropicophyceae uncovered convergent tRNA gene losses and a variant plastid genetic code

**DOI:** 10.1101/530998

**Authors:** Monique Turmel, Adriana Lopes dos Santos, Christian Otis, Roxanne Sergerie, Claude Lemieux

## Abstract

The tiny green algae belonging to the Chloropicophyceae play a key role in marine phytoplankton communities; this newly erected class of prasinophytes comprises two genera (*Chloropicon* and *Chloroparvula*) containing each several species. We sequenced the plastomes and mitogenomes of eight *Chloropicon* and five *Chloroparvula* species to better delineate the phylogenetic affinities of these taxa and to infer the suite of changes that their organelle genomes sustained during evolution. The relationships resolved in organelle-based phylogenomic trees were essentially congruent with previously reported rRNA trees, and similar evolutionary trends but distinct dynamics were identified for the plastome and mitogenome. Although the plastome sustained considerable changes in gene content and order at the time the two genera split, subsequently it remained stable and maintained a very small size. The mitogenome, however, was remodeled more gradually and showed more fluctuation in size, mainly as a result of expansions/contractions of intergenic regions. Remarkably, the plastome and mitogenome lost a common set of three tRNA genes, with the *trnI*(cau) and *trnL*(uaa) losses being accompanied with important variations in codon usage. Unexpectedly, despite the disappearance of *trnI*(cau) from the plastome in the *Chloroparvula* lineage, AUA codons (the codons recognized by this gene product) were detected in certain plastid genes. By comparing the sequences of plastid protein-coding genes from chloropicophycean and phylogenetically diverse chlorophyte algae with those of the corresponding predicted proteins, we discovered that the AUA codon was reassigned from isoleucine to methionine in *Chloroparvula*. This noncanonical genetic code has not previously been uncovered in plastids.

## Introduction

Prasinophytes constitute a paraphyletic assemblage of unicellular, morphologically diversified, and predominantly marine green algae at the base of the Chlorophyta (Leliaert, et al. 2016; Lemieux, et al. 2014; Sym 2015). Apart from the tiny algae belonging to the Mamiellophyceae, which are well known for their important contributions to phytoplankton communities and to primary productivity (Rii, et al. 2016), at least four other prasinophyte lineages (Prasinococcales, Pycnococcaceae, Chloropicophyceae, and Picocystophyceae) comprise species of small size (≤ 5 μm in diameter). A highly reduced, coccoid growth form is thought to confer a distinct advantage to planktonic green algae because the resulting higher surface area to volume ratio enhances the efficiency of nutrient uptake and also because a reduced size helps to escape predators and promotes buoyancy (Grimsley, et al. 2015; Potter, et al. 1997; Sym 2015). Analyses of 18S rRNA metabarcoding data sets have demonstrated that the distribution pattern and habitat of prasinophytes in marine waters vary markedly depending upon the lineage and species examined (Lopes dos Santos, et al. 2017a; Rii, et al. 2016; Tragin and Vaulot 2018). The members of the three genera found in the order Mamiellales (Mamiellophyceae) — *Ostreococcus* (the genus containing the smallest known free-living eukaryote, *O*. *tauri*), *Micromonas* and *Bathycoccus* — are typically found in coastal waters, although some species occur in open oceanic waters (Sym 2015).

More recently, a study based on the analysis of metabarcoding datasets that were produced in the framework of the Tara Oceans expedition and Ocean Sampling Day consortium pointed the key role of the pico/nano-prasinophytes belonging to the lineage sister to the core Chlorophyta (a clade comprising the Chlorodendrophyceae, Pedinophyceae, Trebouxiophyceae, Ulvophyceae and Chlorophyceae (Leliaert, et al. 2016; Lemieux, et al. 2014)) and formerly designated as clade VII (Guillou, et al. 2004) in marine phytoplankton communities, especially in moderately oligotrophic waters (Lopes dos Santos, et al. 2017a; Lopes dos Santos, et al. 2017b). These algae represent the dominant green algal group in open oceanic waters of tropical regions. This finding provided a fresh impetus to investigate the taxonomy and fundamental aspects of the biology of these algae, such as morphology, ultrastructure, and composition of pigments (Lopes dos Santos, et al. 2016; Lopes dos Santos, et al. 2017a; Lopes dos Santos, et al. 2017b). Analysis of the nuclear 18S and plastid 16S rRNA genes from a large sampling of strains and environmental sequences uncovered important diversity within clade VII (Lopes dos Santos, et al. 2017a; Lopes dos Santos, et al. 2017b), leading Lopes dos Santos et al. (2017b) to confer the status of class (Chloropicophyceae) to the lineage. Two major clades that received strong support were each erected to the genus level, *Chloropicon* (clade A) and *Chloroparvula* (clade B), with each genus displaying multiple subclades (A1-A7 and B1-B3) corresponding to different species. However, the precise relationships among the species could not be resolved unambiguously.

Because the chloroplast and mitochondrial genomes, designated hereafter as plastomes and mitogenomes, can be easily sequenced due to their relatively small size and multiple copies/cell, they are often used to infer deep-level phylogenetic relationships among algae and plants and to clarify phylogenetic issues arising from the analysis of a single or few genes (Gitzendanner, et al. 2018; Turmel and Lemieux 2018; Yu, et al. 2018). Moreover, comparative analysis of plastomes and mitogenomes provides the opportunity to gain insights into ancestral architectures and evolution of these genomes. Nine plastomes are currently available for prasinophyte picoalgae (Lemieux, et al. 2014; Moreau, et al. 2012; Robbens, et al. 2007; Turmel, et al. 2009; Worden, et al. 2009). They are smaller in size (< 99 kb), have a reduced gene repertoire (ranging from 100 genes in *Chloropicon primus* to 114 in *Picocystis salinarum*), and are more gene-dense compared to the plastomes of prasinophytes [e.g. *Nephroselmis olivacea* (Turmel, et al. 1999b) and *Pyramimonas parkeae* (Satjarak and Graham 2017; Turmel, et al. 2009)] and core chlorophytes with a larger cell body (Turmel and Lemieux 2018). As revealed by gene mapping on evolutionary trees, the plastome sustained losses of many genes several times independently during the evolution of prasinophyte picoalgae (Turmel and Lemieux 2018). Because the nuclear genomes of mamiellalean picoalgae also feature a reduced size and a high gene density (Grimsley, et al. 2015), it appears that miniaturization of both the nuclear genome and plastome occurred along cell reduction during evolution. The *Chloropicon primus* plastome is currently the sole organelle genome reported for the Chloropicophyceae and at 64.3 kb, it is the smallest plastome documented among photosynthetic green algae (Lemieux, et al. 2014). Unlike the plastomes of the picoalgae *Ostreococcus tauri*, *Micromonas commoda* and *Picocystis salinarum*, it has not retained the large inverted repeat (IR) encoding the rRNA genes, which is commonly found in the majority of green algae and land plants (forming together the Viridiplantae or Chloroplastida) and as observed for other picoalgae, its gene order is substantially scrambled relative to most other prasinophyte plastomes (Lemieux, et al. 2014). Within the Mamiellales order, however, the *Ostreococcus* and *Micromonas* plastomes are highly similar in size and gene content and are essentially colinear.

The mitogenome is usually more divergent than the plastome among green algae. To date, complete or almost complete mitogenome sequences are available for only five prasinophyte picoalgae, which represent the Mamiellales, Prasinococcales and Pycnococcaceae (Moreau, et al. 2012; Pombert, et al. 2013; Robbens, et al. 2007; Turmel, et al. 2010; Worden, et al. 2009). They show important variation in size (24.3 kb in *Pycnococcus provasolii* to 54.5 kb in *Prasinoderma coloniale*) and gene content (36 in *Pycnococcus* to 63 in *Ostreococcus* and *Micromonas*), but no general trend toward a reduced gene complement when they are compared to the mitogenomes of *Nephroselmis* (45.2 kb, 66 genes) (Turmel, et al. 1999a), *Pyramimonas parkeae* (NIES254: 53.4 kb, 58 genes; SCCAP K-0007: 43.3 kb, 59 genes) (Hrda, et al. 2017; Satjarak, et al. 2017) and *Cymbomonas tetramitiformis* (73.5 kb, 56 genes) (Satjarak, et al. 2017). As observed for *Pyramimonas*, the mitogenomes of mamiellalean taxa and *Prasinoderma* carry a large IR with variable gene content. At the level of gene order, mitogenome rearrangements are observed not only across lineages but also within the Mamiellales (Satjarak, et al. 2017).

The goals of the present study were to better delineate the phylogenetic affinities among the different species previously described in the Chloropicophyceae and to gain insights into the evolutionary histories of the plastome and mitogenome within this class. We generated the organelle genome sequences of 13 chloropicophycean taxa and compared the newly sequenced plastomes and mitogenomes to one another and to their counterparts in *Picocystis salinarum* (the unique representative of the Picocystophyceae, a prasinophyte lineage presumed to have emerged just before the Chloropicophyceae). Our phylogenomic analyses produced a robust phylogeny on which the structural changes inferred from genome comparisons were mapped. For each organelle genome, important differences were observed between the *Chloropicon* and *Chloroparvula* genera as well as between each genus and *Picocystis* or any other previously examined prasinophyte. Most unexpected was our discovery of a non-canonical genetic code in the *Chloroparvula* plastome.

## Materials and Methods

### Strains and culture conditions

Strains of *Chloropicon primus* (CCMP 1205) and *Picocystis salinarum* (CCMP 1897) were obtained from the National Center for Marine Algae and Microbiota (NCMA, https://ncma.bigelow.org) and were maintained in K medium at 18°C. The 12 other chloropicophycean strains originated from the Roscoff Culture Collection (RCC, http://roscoff-culture-collection.org) and were maintained in L1 medium at 22°C. All algal cultures were grown synchronously under 12-hour light/dark cycles.

### Genome sequencing and annotation

The *Chloropicon primus* and *Picocystis* mitogenomes were assembled in the course of sequencing an A+T-rich DNA fraction that also contained the plastome; the methodologies for sequencing these algal mitogenomes were therefore the same as previously described for their plastomes (Lemieux, et al. 2014). For the other chloropicophycean strains, the procedure used was as follows. Total cellular DNA was extracted using the EZNA HP Plant DNA Mini kit of Omega Bio-tek (Norcross, GA, USA). Paired-end libraries (inserts of 500 bp) were prepared using the Illumina TruSeq DNA kit and DNA was sequenced on the MiSeq platform (300 cycles). Both library construction and DNA sequencing were performed at the Plateforme d’Analyses Génomiques of Université Laval (Québec, QC, Canada) (http://www.ibis.ulaval.ca/services/analyse-genomique/). After removing low-quality bases and adapters using AfterQC (Chen, et al. 2017), reads were assembled using Spades 3.11.1 (Bankevich, et al. 2012). Contigs of plastid and mitochondrial origins were identified by BlastN and BlastX searches against a local database of green plant organelle genomes and then were scaffolded in Sequencher 5.4.6 (Gene Codes Corporation, Ann Arbor, MI, USA).

Plastome and mitogenome annotations were performed using a custom-built suite of bioinformatics tools as previously described (Turmel, et al. 2017). Genes encoding tRNAs were identified independently using tRNAscan-SE (Lowe and Eddy 1997). Circular maps of organelle genomes were drawn with OGDraw (Lohse, et al. 2013).

### Phylogenomic analyses

Phylogenomic trees were inferred from plastome and mitogenome datasets that each comprised 14 taxa (chloropicophyceans + *Picocystis*). The plastome dataset (PCG12RNA) was prepared from the first and second codon positions of 71 protein-coding genes and the sequences coding for 3 rRNAs and 28 tRNAs, while the mitogenome dataset (also designated as PCG12RNA) was derived from the first and second codon positions of 36 protein-coding genes and the sequences coding for 2 rRNAs and 26 tRNAs. Following alignment of the deduced amino acid sequences from the individual protein-coding genes using Muscle 3.7 (Edgar 2004) and conversion of the resulting alignments to codon alignments, poorly aligned and divergent regions in each gene alignment were excluded using Gblocks v0.91b (Castresana 2000) with the -t=c, -b3=5, -b4=5 and -b5=half options, and individual alignments were concatenated using Phyutility v2.2.6 (Smith and Dunn 2008). The third codon positions of the resulting alignment were excluded using Mesquite v3.51 (Maddison and Maddison 2018) to produce the PCG12 data set, which was then merged with the concatenated alignment of the RNA-coding genes to produce the PCG12RNA data set. To generate this concatenated alignment, individual genes were aligned using Muscle 3.7 (Edgar 2004), ambiguously aligned regions in each gene alignment were removed using TrimAl v1.4 (Capella-Gutierrez, et al. 2009) with the options block=6, gt=0.9, st=0.4 and sw=3, and alignments were concatenated using Phyutility v2.2.6 (Smith and Dunn 2008).

The plastome and mitogenome PCG12RNA data sets were analyzed using maximum likelihood (ML) and Bayesian methods of phylogenetic reconstruction. ML analyses were performed using RAxML v8.2.12 (Stamatakis 2014) and the GTR+Γ4 model of sequence evolution, with each of the data sets being partitioned into gene groups and the model applied to each partition. The partitions included two RNA gene groups (rRNA and tRNA genes) in addition to the protein-coding gene partitions. Confidence of branch points was estimated by bootstrap analysis with 100 replicates. Bayesian analyses were performed using PhyloBayes 4.1 (Lartillot, et al. 2009) under the heterogeneous CAT-GTR model. Two independent chains were run for 5,000 cycles and consensus topologies were calculated from the saved trees using the BPCOMP program of PhyloBayes after a burn-in of 1,000 cycles. Under these conditions, the largest discrepancy value observed across all bipartitions in the consensus topologies (maxdiff) was lower than 0.10, indicating that convergence between the chains was achieved.

In addition, a separate plastome-based phylogenomic tree was inferred using an amino acid dataset comprising 167 green plant taxa. This dataset, which was derived from 79 protein-coding genes, was assembled and analyzed under ML as described previously (Turmel, et al. 2017), except that IQ-Tree 1.6.7 (Nguyen, et al. 2015) and ultrafast bootstrap approximation (Hoang, et al. 2018) were used for the tree reconstruction and bootstrap analysis, respectively. Taxon sampling included the 129 green plants that were recently examined in the phylogenomic analysis reported by Turmel and Lemieux (2018); the GenBank accessions of the plastomes for the 38 additional taxa are as follows: *Acorus calamus*, NC_007407; *Anthoceros formosae*, NC_004543; *Arabidopsis thaliana*, NC_000932; *Carteria crucifera*, KT624870-KT624932; *Chlamydomonas asymmetrica*, KT624933-KT625007; *Chlamydomonas moewusii*, EF587443-EF587503; *Chlorogonium capillatum*, KT625085-KT625091; *Chloromonas radiata*, KT625008-KT625084; *Chloromonas typhlos*, KT624630-KT624716; *Golenkinia longispicula*, KT625092-KT625150; *Haematococcus lacustris*, KT625205-KT625298; *Hydrodictyon reticulatum*, NC_034655; *Huperzia lucidula*, NC_006861; *Lobochlamys culleus* (KT625151-KT625204; *Lobochlamys segnis*, KT624806-KT624869; *Marchantia polymorpha*, NC_001319; *Microglena monadina*, KT624717-KT624805; *Nicotiana tabacum*, NC_001879; *Oryza sativa*, NC_001320; *Pediastrum duplex*, NC_034654; *Physcomitrella patens*, NC_005087; *Pinus thunbergii*, NC_001631; *Pyramimonas parkeae*, KX013546; *Stephanosphaera pluvialis*, KT625299-KT625409; *Syntrichia ruralis*, NC_012052; *Trebouxia aggregata*, EU123962-EU124002.

### Gene order analyses

Genome-scale sequence alignments were carried out using LAST 9.5.6 (Frith, et al. 2010) and analyses of genome rearrangements were performed using the ProgressiveMauve algorithm of Mauve 2.3.1 (Darling, et al. 2010). The number of reversals separating the plastomes and mitogenomes were estimated using GRIMM v2.01 (Tesler 2002) and gene order matrices containing the genes shared by all these genomes (plastomes, 95 genes; mitogenomes, 53 genes). The gene order matrices were also used to infer ML trees using the phylogeny reconstruction option of MLGO (Hu, et al. 2014) and confidence of branch points was estimated by 1,000 bootstrap replications. The user tree option of MGR v2.03 (Bourque and Pevzner 2002) was used to infer the number of gene reversals on each branch of the topology recovered from phylogenomic analyses.

### Codon usage and tRNA analyses

Analysis of codon usage in individual organelle genome was carried out using CUSP in EMBOSS 6.6.0 (Rice, et al. 2000) and concatenated sequences of protein-coding genes. CUSP was also used to identify the subset of *Chloroparvula* plastid genes featuring AUA codons. The positions of these AUA codons as well as the corresponding codons and associated amino acid(s) in green algal orthologs were identified from gene alignments using the “translate as cDNA” option of Jalview (Waterhouse, et al. 2009), which dynamically links codon alignments with predicted proteins in a split-frame view. The same program was employed to identify the codons corresponding to universally conserved isoleucine residues in plastome-encoded proteins of phylogenetically diverse green algae.

To determine in which part of the tRNA secondary structure fall the nucleotide differences observed between the *Chloroparvula* and *Chloropicon* elongator tRNA^Met^(CAU) sequences, secondary structures of these tRNAs were determined using tRNAscan-SE (Lowe and Eddy 1997) and a consensus secondary structure was drawn manually.

## Results

Of the 13 strains sampled from Chloropicophyceae (table 1), eight belong to the *Chloropicon* genus; these represent all seven clades identified in this genus and include two distinct isolates of *Chloropicon roscoffensis* (A4), one from the English Channel (RCC1871) and the other from the Pacific Ocean (RCC2335). The remaining strains represent four separate clades of the *Chloroparvula* genus (B1-B3 and B) and include two B2-clade members from the Indian (RCC999) and Pacific (RCC696) Oceans. This sampling strategy allowed us to track organelle genome changes on both macro- and micro-evolutionary scales. In addition, the mitogenome sequence of *Picocystis salinarum* (Picocystophyceae) was generated in order to compare the newly sequenced chloropicophycean mitogenomes with a close relative from a distinct prasinophyte class. Although all chloropicophycean taxa, except *Chloropicon primus*, were not available as axenic cultures, their entire plastomes and mitogenomes could be assembled with no ambiguity as circular-mapping molecules (supplementary figs. S1 and S2, Supplementary Material online). Considering that an accurate phylogenetic framework is essential to track the suite of genomic changes that took place during chloropicophycean evolution, we will present our phylogenomic analyses before elaborating further on the structural features of the newly sequenced organelle genomes.

**Table 1.**
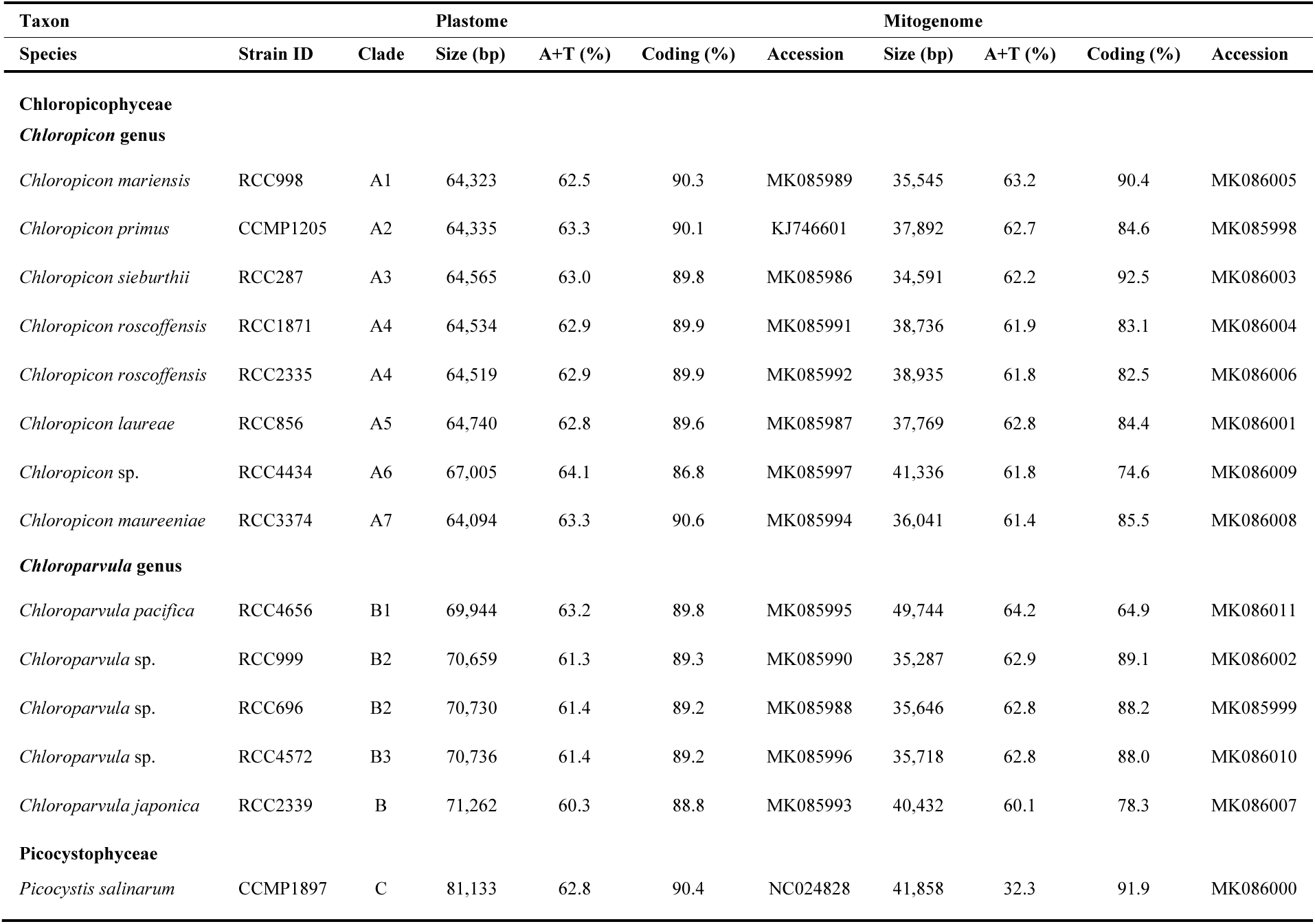
Characteristics of plastomes and mitogenomes from chloropicophycean taxa and *Picocystis salinarum*.

### Plastome- and Mitogenome-Based Phylogenomic Analyses

Plastome and mitogenome-based phylogenomic trees were inferred from 102 and 64 concatenated genes, respectively, using ML and Bayesian methods and *Picocystis salinarum* as outgroup (fig. 1). Both trees display exactly the same topology and their branches are well resolved, leaving little ambiguity regarding the relative positions of the sampled chloropicophycean taxa. Overall, the relationships are consistent with the trees previously inferred from nuclear 18S and plastid 16S rRNA genes (Lopes dos Santos, et al. 2017b); however, there are differences in the relative positions of the *Chloroparvula* sp. B2 and B3 strains and also of *Chloropicon mariensis* (A5) and *Chloropicon laureae* (A1). In addition, the position of *Chloroparvula japonica* (B) relative to other clades within this genus, which received poor support in the tree based on the concatenated rRNA genes, is supported by maximal bootstrap and Bayesian posterior values in the phylogenomic trees.

**Fig. 1.**
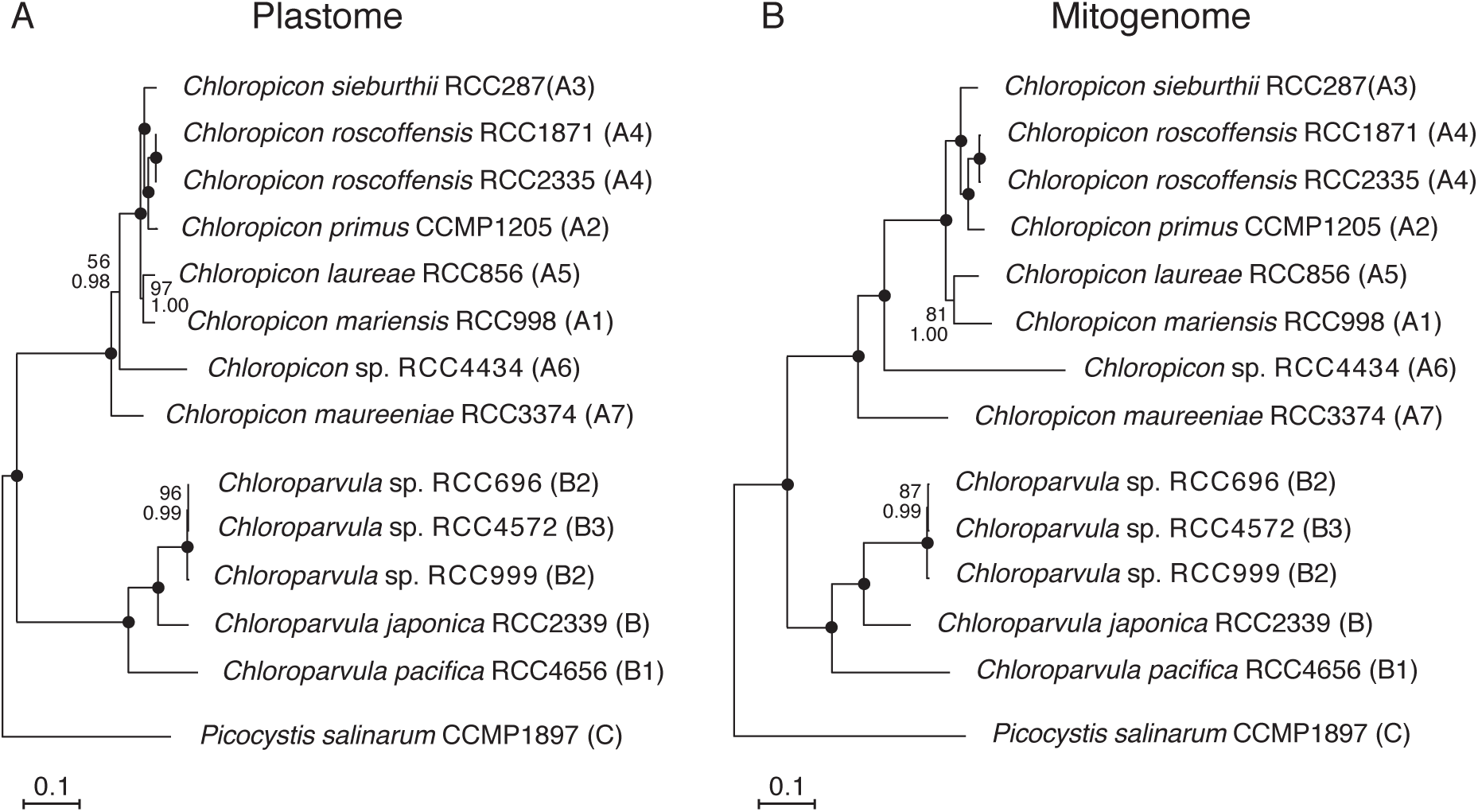
Organelle phylogenomic trees showing the relationships among chloropicophycean taxa. (*A*) Plastome-based phylogeny inferred from a data set of 102 genes (first and second positions of 71 protein-coding genes, 3 rRNA genes, and 28 tRNA genes). (*B*) Mitogenome-based phylogeny inferred from a data set of 64 genes (first and second positions of 36 protein-coding genes, 2 rRNA genes, and 26 tRNA genes). The trees shown here are the best-scoring ML trees that were inferred under the GTR+Γ4 model. Support values are reported on the nodes: from top to bottom are shown the bootstrap support (BS) values for the RAxML GTR+Γ4 analyses and the posterior probability (PP) values for the PhyloBayes CATGTR+Γ4 analyses. Black dots indicate that the corresponding branches received BS and PP values of 100%.

It is worth noting that the internal nodes in the *Chloropicon* lineage are separated by greater distances in the mitogenome tree (fig. 1B) than in the plastome tree (fig. 1A), suggesting different rates of evolution for these organelle genomes. Similarly, the internal branch in the *Chloroparvula* lineage that connects the common ancestor of the B2-B3 strains is longer in the mitogenome tree.

Given that taxon sampling may influence the branching order of lineages, we tested whether the Chloropicophyceae remain sister to all core chlorophytes when plastome-based trees are constructed using completely sequenced green algal plastomes available in public databases, including those newly sampled in this study. A ML tree was inferred from an amino acid dataset derived from 79 genes of 167 green plant taxa representing various lineages of the Chlorophyta and Streptophyta (fig. 2). This tree identified the Chloropicophyceae as sister to all core chlorophytes, with the Picocystophyceae + Pycnococcaceae clade emerging as the preceding lineage. Judging from the depth of the positions occupied by the common ancestor of all *Chloropicon* and by that of all *Chloroparvula* species, each of these genera exhibits important genetic diversity and one notes that these divergence levels far exceed that observed for the combined *Micromonas* and *Ostreococcus* genera, which together with *Bathycoccus* constitute the Mamiellales order of the Mamiellophyceae.

**Fig. 2.**
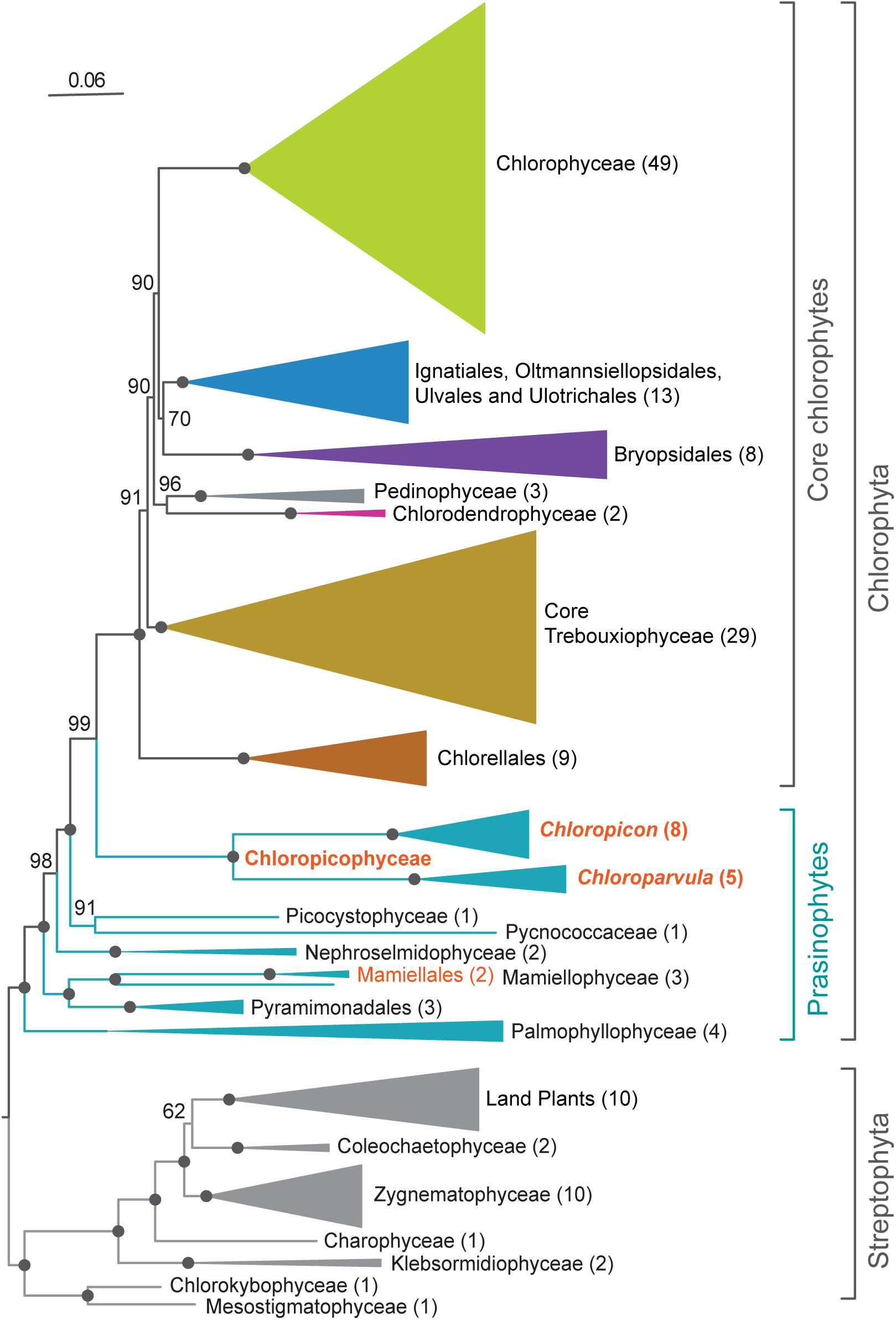
Plastome-based phylogenomic tree showing the position of the Chloropicophyceae relative to other classes and major lineages of the Chlorophyta. This ML tree was inferred from a data set of 79 proteins from 167 green plants using IQ-Tree under the GTR+R4 model. Bootstrap support values are reported on the nodes. The clades that received maximal support were collapsed and represented as triangles with sizes proportional to the number of taxa (indicated in parentheses). Black dots indicate that the corresponding branches received 100% bootstrap support.

### Plastome Organization

Unlike the 81.1-kb plastome of *Picocystis*, which contains 114 genes, all chloropicophycean plastomes lack an IR encoding the rRNA genes and also differ by their substantially smaller size and gene repertoire as well as by the absence of introns (table 1, fig. 3A and supplementary fig. S1, Supplementary Material online). At 64,094 bp, the *Chloropicon maureeniae* (A7) plastome exhibits the smallest size among chloropicophycean plastomes and is also the smallest known among photosynthetic green algae. Plastomes of *Chloroparvula* species are about 7 kb larger but contain fewer genes (102 versus 97) in comparison to *Chloropicon* species. Within each of these genera, however, there is very little variation in plastome size, gene content and density of coding sequences (table 1 and fig. 3A). Given that a few genes in the *Chloroparvula* plastomes, such as the highly variable *ycf1* encoding a putative subunit of the plastid protein import apparatus, the TOC/TIC machinery (Kikuchi, et al. 2013), exhibit multiple insertions and are expanded relative to the corresponding *Chloropicon* genes (supplementary fig. S3, Supplementary Material online), all examined chloropicophycean plastomes share similar gene densities.

**Fig. 3.**
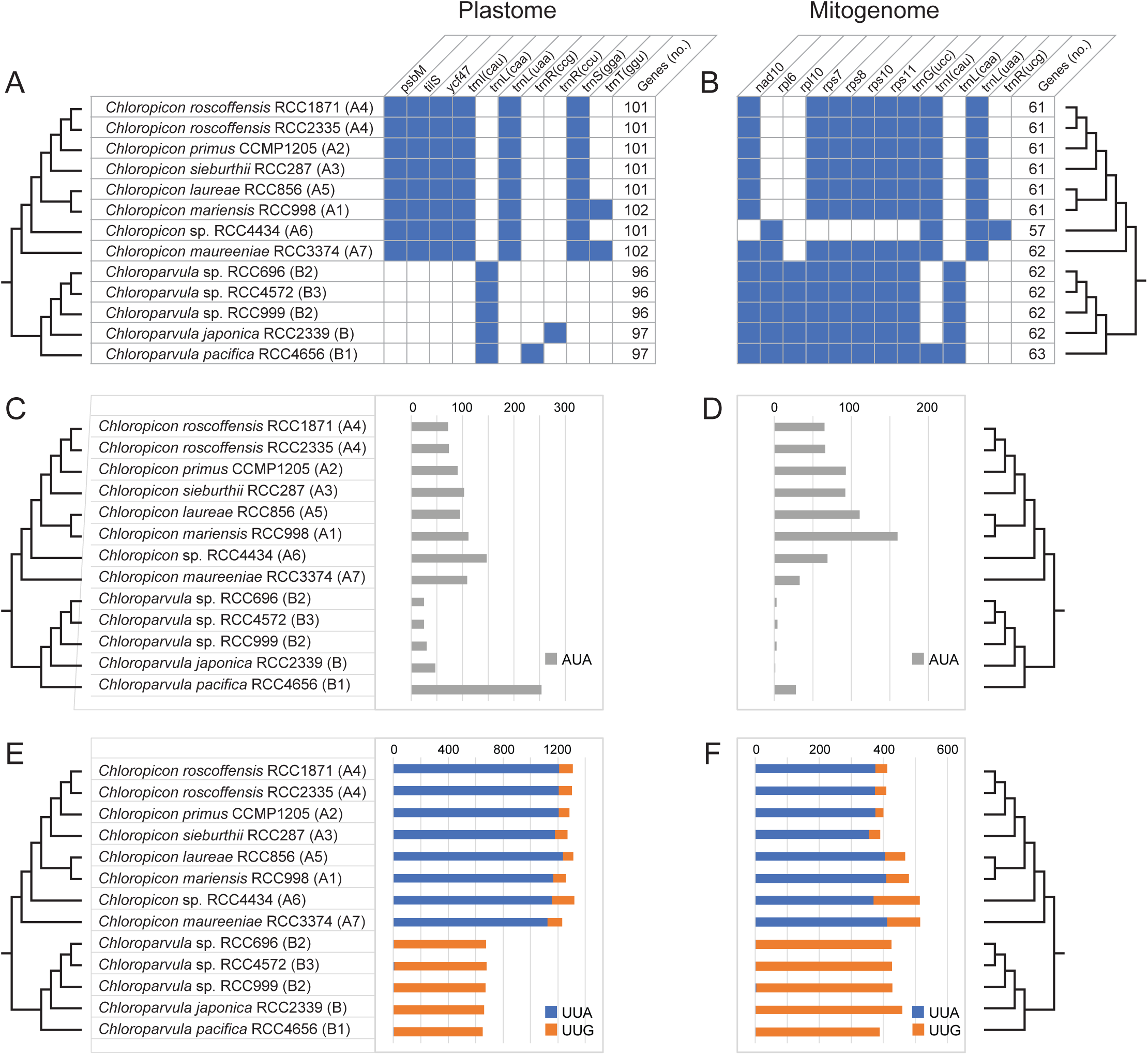
Changes in gene content and codon frequency among chloropicophycean plastomes and mitogenomes. (*A, B*) Distribution of variable genes and total number of genes. The presence of a gene is denoted by a blue box. All plastomes share the following 95 genes: *accD, atpA,B,E,F,H,I, chlB,L,N, clpP, ftsH, infA, minD, petA,B,D,G,L, psaA,B,C,J, psbA,B,C,D,E,F,H,I,J,K,L,N,T,Z, rbcL, rpl2,5,14,16,19,20,23,32,36, rpoA,B,C1,C2, rps2,3,4,7,8,9,11,12,14,18,19, rrf, rrl, rrs, tufA, ycf1,3,4,12,20, trnA*(ugc), *C*(gca), *D*(guc), *E*(uuc), *F*(gaa), *G*(gcc), *G*(ucc), *H*(gug), *I*(gau), *K*(uuu), *L*(uag), *Me*(cau), *Mf*(cau), *N*(guu), *P*(ugg), *Q*(uug), *R*(acg), *R*(ucu), *S*(gcu), *S*(uga), *T*(ugu), *V*(uac), *W*(cca), *Y*(gua). All mitogenomes share the following 53 genes: *atp1,4,6,8,9, cob, cox1,2,3, mttB, nad1,2,3,4,4L,5,6,7,9, rpl5,14,16, rps2,3,4,12,13,14,19, rnl, rns, trnA*(ugc), *C*(gca), *D*(guc), *E*(uuc), *F*(gaa), *G*(gcc), *H*(gug), *I*(gau), *K*(uuu), L(uag), *Me*(cau), *Mf*(cau), *N*(guu), *P*(ugg), *Q*(uug), *R*(acg), *R*(ucu), *S*(gcu), *S*(uga), *V*(uac), *W*(cca), *Y*(gua). Duplicated gene copies, including those observed for the *Chloroparvula pacifica* plastid *trnP*(ugg) and *trnW*(cca) genes, were counted once. (*C, D*) Changes in frequency of the AUA codon associated with the presence/absence of *trnI*(cau). (*E, F*) Changes in frequencies of the UUA and UUG codons associated with the presence/absence of *trnL*(uaa) and *trnL*(caa), respectively.

Ten genes account for the differences in coding capacities of chloropicophycean plastomes and of these, seven are genus-specific (fig. 3A). Three protein-coding genes, including *tilS*, as well as three tRNA genes [*trnI*(cau), *trnL*(uaa), and *trnS*(gaa)] are specific to *Chloropicon*, whereas only the *trnL*(caa) gene is specific to *Chloroparvula*. The fact that both *trnI*(cau) and *tilS* are either present or missing is not a coincidence, as *tilS* encodes the tRNA(Ile)-lysidine synthase that is required to convert to lysidine the cytidine found at the wobble position of the AUA codon-specific tRNA^Ile^(CAU); this conversion changes the amino acid specificity of the tRNA from methionine to isoleucine (Soma, et al. 2003).

Given that *trnI*(cau) occurs in all *Chloropicon* taxa but is absent in all *Chloroparvula* taxa, one would predict the AUA codon to be less frequent in *Chloroparvula* than in *Chloropicon* plastomes or to be even missing entirely. This prediction, which is based on the assumption that no nuclear-encoded tRNA^Ile^(CAU) is targeted to the plastid, was verified by measuring the number of occurrences of the AUA codon in the protein-coding genes of each chloropicophycean plastome. It was found that all five *Chloroparvula* species, with the exception of *Chloroparvula pacifica*, exhibit much fewer AUA codons compared to the *Chloropicon* species (fig. 3C and supplementary table S1, Supplementary Material online). In which genes are located the AUA codons in *Chloroparvula*? When their positions correspond to universally or almost universally conserved amino acid positions in proteins of phylogenetically diverse green algae, do they generally specify isoleucine or a different amino acid residue? To address these questions, the protein-coding genes of *Chloroparvula japonica* (RCC2339) and *Chloroparvula pacifica* (RCC4656) carrying AUA codons were aligned with their orthologs in other green algae and the correspondence between these AUA codons and specific amino acid residues was visualized with the aid of Jalview (Waterhouse, et al. 2009). Most of the *Chloroparvula* genes containing AUA codons encode proteins that are not directly implicated in photosynthesis (e.g. *chlB* and *chlN* encode proteins required for light-independent chlorophyll synthesis) (fig. 4), and to our surprise, the AUA codons occupying universally conserved positions at the protein level were found to invariably correspond to AUG codons in *Chloropicon* species as well as in the other green algae examined (fig. 5). As expected, the AUA codons present at universally conserved amino acid positions in the latter algae are decoded as isoleucine residues (supplementary fig S4, Supplementary Material online). These results suggest that in *Chloroparvula* plastomes, the AUA codon is decoded as methionine and that two codons (AUU and AUC) instead of three, are read as isoleucine. Considering that RNA editing, a process mainly responsible mainly for U→C and C→U changes in organelle transcripts of land plants, has not been detected in green algae (Cahoon, et al. 2017), conversion of the AUA codon to AUG at the RNA level is very unlikely to be responsible for decoding of AUA as methionine in *Chloroparvula*. To explore the molecular basis for the reassignment of AUA to methionine, the *Chloroparvula* elongator tRNA^Met^(CAU) gene sequences were aligned with the corresponding sequences of *Chloropicon* and other green algae and specific mutations were found in the *Chloroparvula* sequences (fig. 6).

**Fig. 4.**
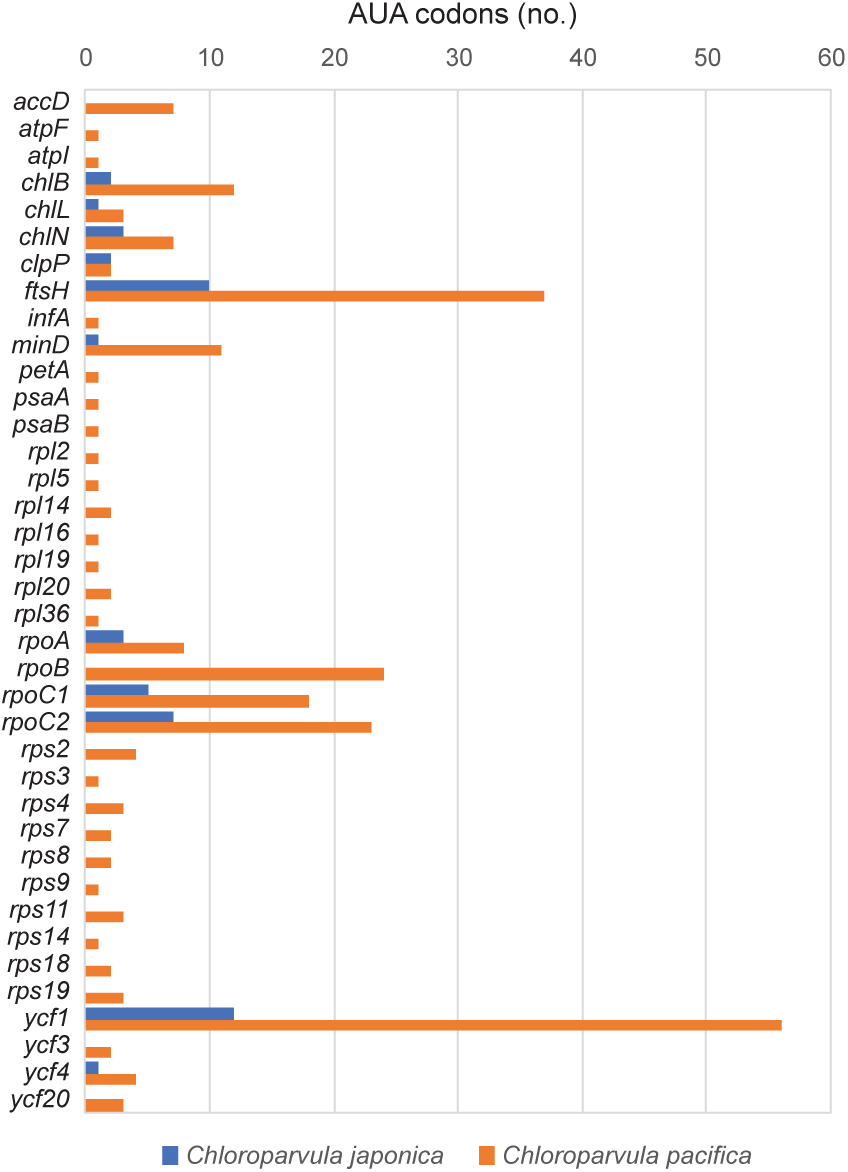
Number of AUA codons in plastid genes of *Chloroparvula japonica* and *Chloroparvula pacifica*.

**Fig. 5.**
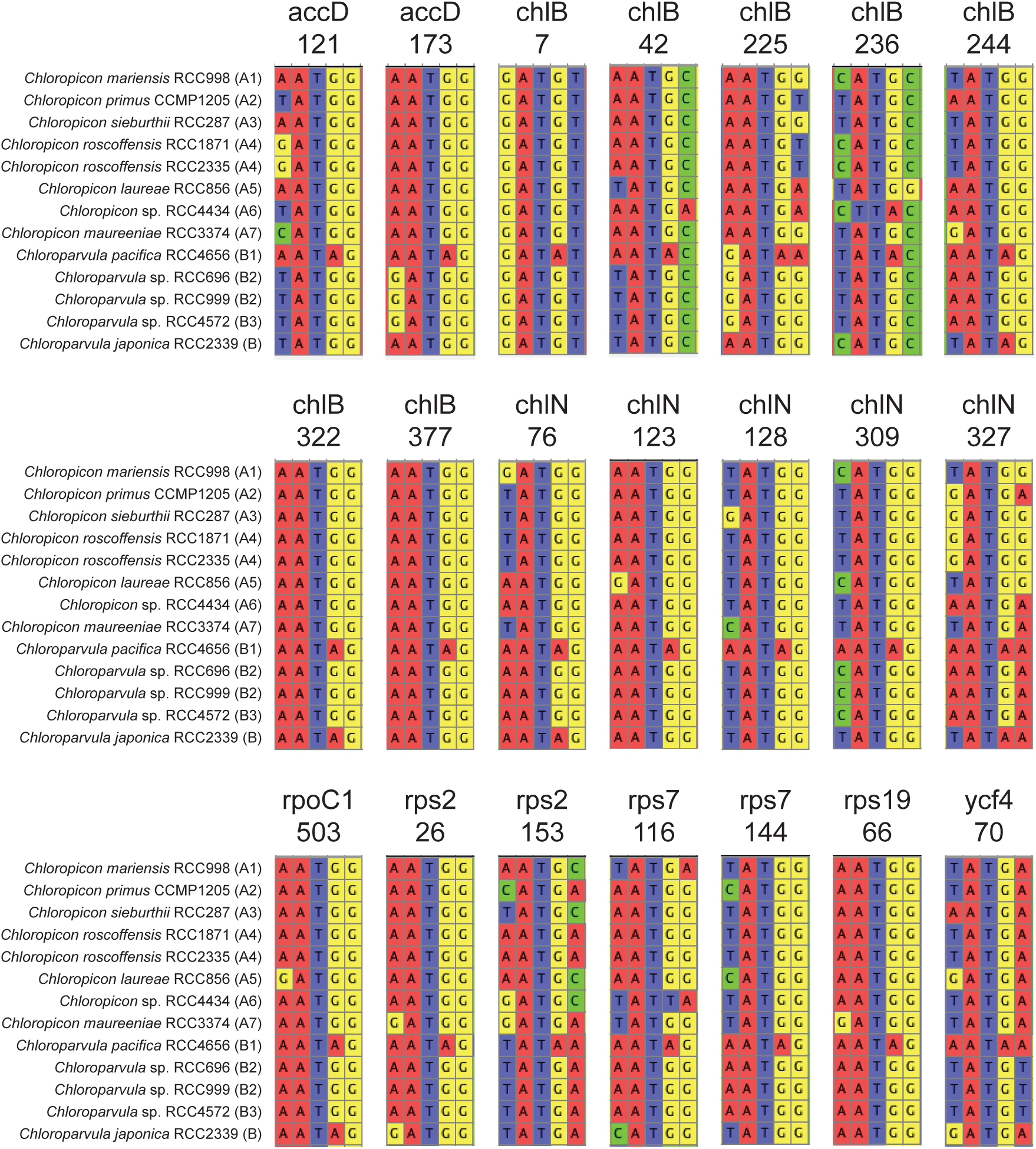
Partial gene alignments including AUA codons from *Chloroparvula* plastomes. These codons fall within universally or almost universally conserved sites at the protein level and correspond to AUG codons in orthologous plastid genes of *Chloropicon* and other green algae. The 21 sites containing AUA codons are distributed among eight protein-coding genes; the number below the gene name indicates the position of the AUA codon in the *Chloroparvula pacifica* gene.

**Fig. 6.**
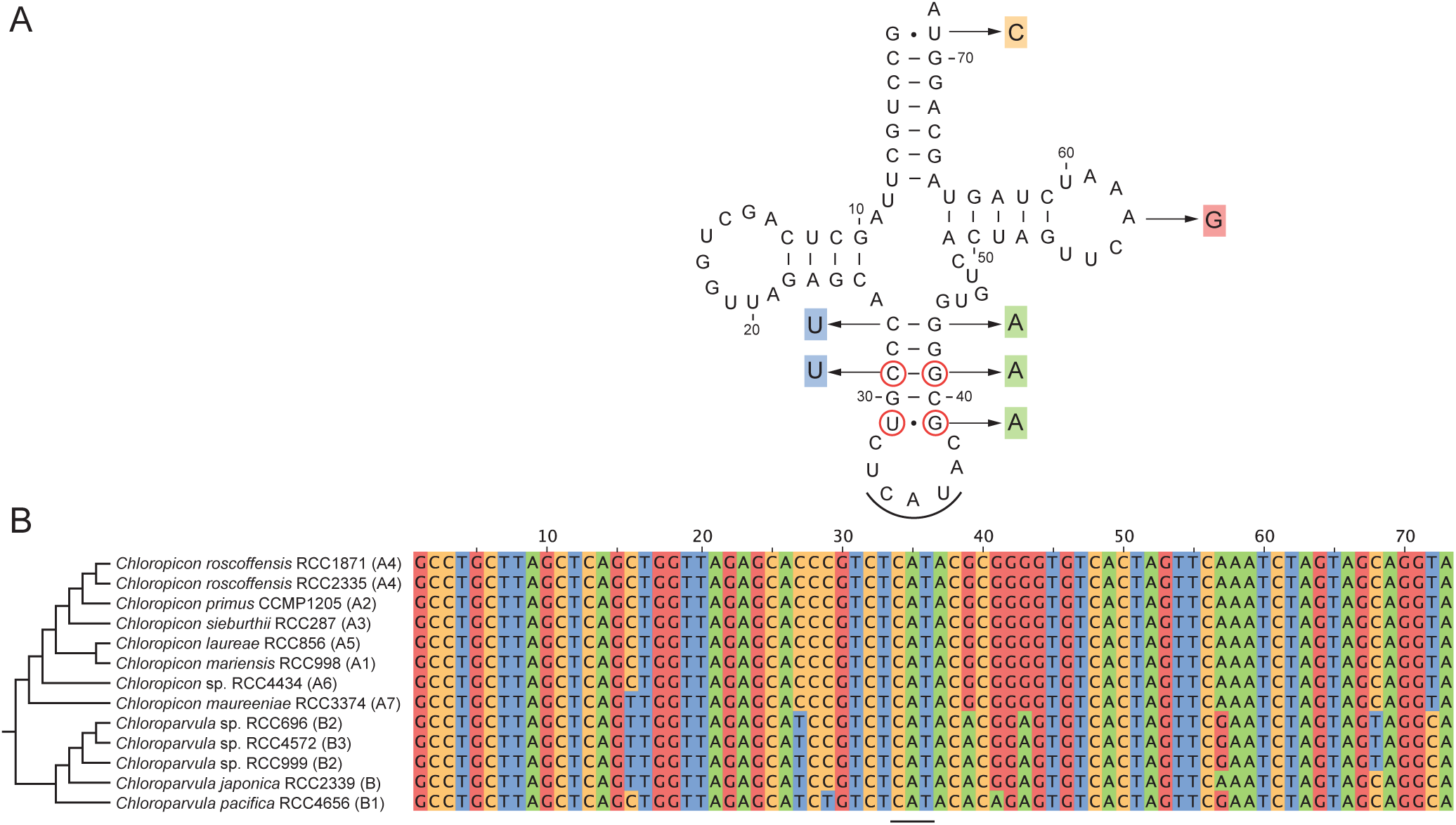
Evolution of the plastid elongator tRNA^Met^(CAU) in the Chloropicophyceae. (*A*) Consensus secondary structure model of the *Chloropicon* elongator tRNA^Met^(CAU) showing the variant nucleotides observed in *Chloroparvula pacifica*. The circled nucleotides in the anticodon stem are conserved in chlorophyte tRNAs^Met^(CAU). (*B*) Alignment of *trnMe*(cau) gene sequences from all examined chloropicophycean taxa. The anticodon sequence is underlined.

With regards to the signature *trnL*(uaa) and *trnL*(caa) genes, one also expects to find highly different frequencies for the UUA and UUG codons in *Chloropicon* and *Chloroparvula* plastomes, given that the tRNA^leu^(UAA) recognizes both the UUA and UUG codons (the third position of the latter codon interacting through wobble pairing) and that the tRNA^leu^(CAA) recognizes solely the UUG codon. As predicted, the UUA codon was found to be prevalent in *Chloropicon* taxa although the UUG codon was also detected, and contrasting with this observation, the UUG codon was identified almost exclusively in *Chloroparvula* species (fig. 3E and supplementary table S1, Supplementary Material online). In the case of *trnS*(gga), its retention in *Chloropicon* and absence in *Chloroparvula* have no major impact on codon frequency because the tRNA encoded by this gene has a redundant role compared to that encoded by the universally present *trnS*(uga), which can recognize all four UCN codons through superwobbling (Alkatib, et al. 2012). Similarly, the three tRNA species encoded by genes present in only one or two taxa [tRNA^thr^(GGU), tRNA^arg^(CCG), and tRNA^arg^(CCU) as shown in fig. 3A] are redundant with the products of *trnT*(ugu), *trnR*(acg), and *trnR(*ucu), which recognize four, four, and two codons, respectively, through superwobbling or wobbling (Alkatib, et al. 2012). It was found that both *trnT*(ggu) and *trnR*(ccg) have several orthologs in other chlorophyte plastomes; however, Blast searches of the NCBI database using *trnR*(ccu) identified strong hits only to *trnR*(ccg) genes from two isolates of the prasinophyte *Pyramimonas parkeae*, suggesting that the latter tRNA gene arose from duplication of *trnR*(ucu) and subsequent mutation of the anticodon.

The level of synteny among chloropicophycean plastomes was analyzed using Mauve (fig. 7A). The *Picocystis* plastome was not included in this analysis because it is extremely rearranged compared to these genomes. The *Chloropicon* plastomes were found to differ considerably from the *Chloroparvula* plastomes at the gene organizational level; however, an identical gene order was observed for all strains belonging to the same genus, with the exception of *Chloropicon sp.* RCC4434 (A6) and *Chloroparvula pacifica*. The number of gene reversals separating the rearranged plastomes was estimated using GRIMM (supplementary fig. S5A, Supplementary Material online) and MLGO (fig. 7C); in both analyses, the input data was a matrix of signed gene order comprising the 95 genes shared by all taxa, with the output being a distance matrix of the estimated numbers of reversals for all pairwise comparisons in the case of GRIMM and a tree generated with this matrix in the case of MLGO. GRIMM identified more than 50 reversals between the *Chloropicon* and *Chloroparvula* plastomes, but only two reversals between the *Chloropicon sp.* RCC4434 (A6) and all other *Chloropicon* plastomes and 19 reversals between the *Chloroparvula pacifica* and all other *Chloroparvula* plastomes. The results obtained with MLGO were consistent with these results.

**Fig. 7.**
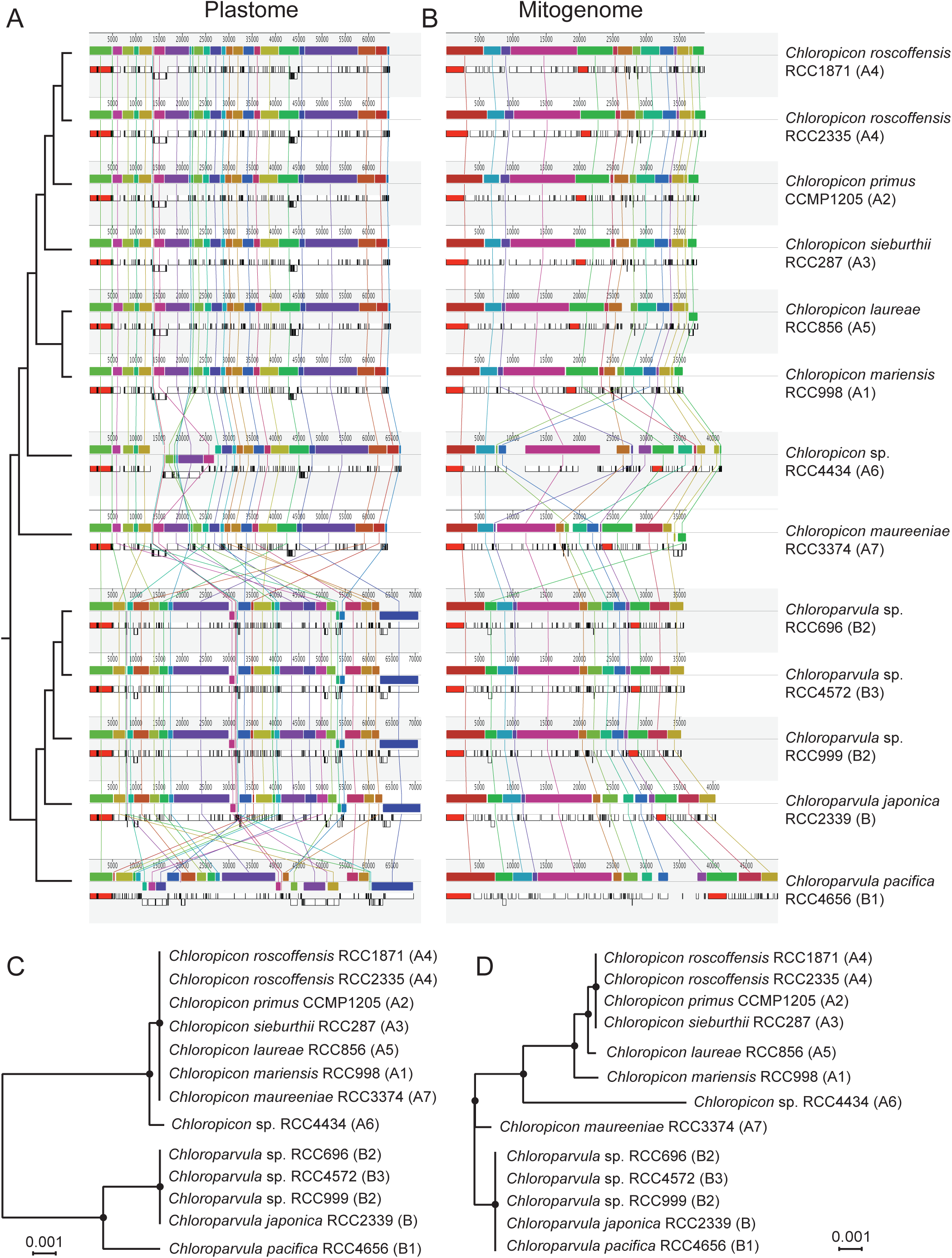
Extent of gene rearrangements in chloropicophycean plastomes and mitogenomes. (*A, B*) Analysis of gene order using Mauve 2.3.1. Locally collinear blocks of genome sequences are represented by boxes of identical color and similarly colored blocks are connected by lines. Blocks above the center line of the aligned regions are in the same orientation as in the reference (RCC1871) genome sequence, while those below this line are in the reverse orientation. (*C, D*) Phylogenetic relationships inferred from gene order datasets using the tree reconstruction option of MLGO. The plastome and mitogenome datasets included 95 and 53 genes, respectively. Consensus trees of 1000 bootstrap replicates are shown; black dots indicate that the corresponding branches received 100% bootstrap support.

Although nearly all plastomes from the same genus were colinear in nucleotide sequence, only those from the most closely related taxa (the *Chloropicon roscoffensis* taxa and the B2/B3 *Chloroparvula* sp. strains) could be aligned over their entire length using LAST. The *Chloroparvula* sp. RCC696 and RCC4572 revealed the lowest density of single nucleotide polymorphisms (SNPs), followed closely by the two *Chloropicon roscoffensis* strains (fig. 8 and supplementary table S2, Supplementary Material online).

**Fig. 8.**
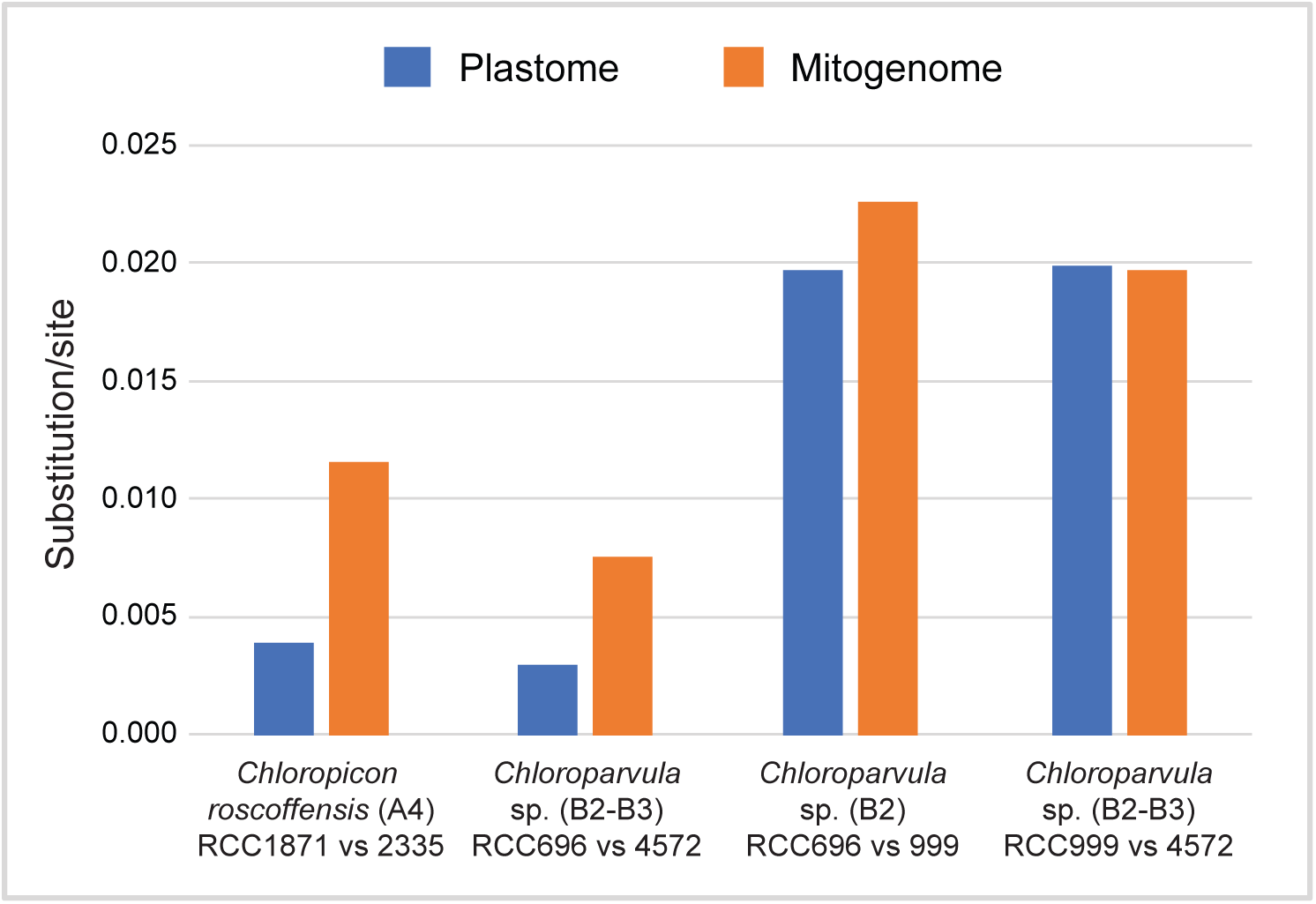
Relative numbers of substitutions per site in the plastomes and mitogenomes of *Chloropicon roscoffensis* and *Chloroparvula* sp. (B2 and B3) strains. To obtain these estimates, whole plastomes and mitogenomes from pairs of closely related strains were aligned and SNPs were identified in the aligned sequences.

### Mitogenome Organization

The 41.9-kb *Picocystis* mitogenome displays important differences relative to its counterparts in the Chloropicophyceae; notably, it features a very scrambled gene order relative to the latter genomes, an unusually low A+T content (table 1), and a 6.3-kb IR containing the *rns*, *rnl*, *trnMe*(cau) and *cob* genes (supplementary fig. S6, Supplementary Material online). Although the number of encoded genes is practically the same in all chloropicophycean mitogenomes, noticeable variations are seen at the genome size and gene density levels within each of the two genera (table 1). The *Chloropicon pacifica* mitogenome is the largest (49.7 kb) and least compact (64.9% coding sequence), followed by the *Chloropicon* sp. RCC4434 mitogenome, which has 57 genes instead of 61 or 62 as in other chloropicophyceans. Whereas no intron is present in the *Picocystis* mitogenome, a group I intron encoding a LAGLIDADG endonuclease occurs at site 1931 of the large subunit rRNA gene (*rnl*) in *Chloropicon sieburthii*, *Chloropicon roscoffensis* RCC2335, *Chloropicon laureae*, and *Chloroparvula pacifica*; moreover, in the latter strain, another group I intron also encoding a LAGLIDADG endonuclease is found at site 793 of the small subunit rRNA gene (*rns*) (supplementary fig. S2, Supplementary Material online).

Three genes distinguish the *Chloropicon* and *Chloroparvula* genera: *trnL*(uaa) is specific to *Chloropicon*, whereas the ribosomal protein-coding gene *rpl10* and *trnL*(caa) are specific to *Chloroparvula* (fig. 3B). Therefore, as reported above for the plastome, *trnL*(uaa) and *trnL*(caa) are mutually exclusive genes in the mitogenome, and the distribution patterns of the UUA and UUG codons in *Chloropicon* and *Chloroparvula* taxa are in complete agreement with those observed for the plastome-encoded genes (fig. 3F and supplementary table S1, Supplementary Material online). The *rpl6* gene, which is present in all *Chloroparvula* strains, was identified in the *Chloropicon* genus but only in the representative of the earliest-diverging lineages (*Chloropicon maureeniae and Chloropicon sp*. RCC4434); similarly, *trnI*(cau), a gene present in all *Chloropicon* taxa, was retained solely in the strain representing the earliest branch of the *Chloroparvula* genus (*Chloroparvula pacifica*). In agreement with the distribution of the latter gene, the AUA codon recognized by tRNA^ile^(CAU) is abundant in *Chloropicon* strains and in *Chloroparvula pacifica* but was detected at fewer than four sites in other *Chloroparvula* strains (fig. 3D and supplementary table S1, Supplementary Material online). The *trnR*(ucg) gene was identified solely in *Chloropicon* sp. RCC4434; this gene is rarely found in green algal mitogenomes and Blast searches of the NCBI database using the *Chloropicon* sp. RCC4434 sequence identified *trnR*(acg) from several chlorophytes as best hits, suggesting that it was possibly derived by duplication of *trnR*(acg) followed by mutation of the anticodon.

While the mitogenomes of all examined *Chloroparvula* strains are colinear, differences in gene order are observed for a few strains sampled from the *Chloropicon* lineage (figs. 7B and 7D, and supplementary fig. S5B, Supplementary Material online). In particular the representatives of the two earliest-diverging *Chloropicon* lineages (*Chloropicon* sp. RCC4434 and *Chloropicon maureeniae*), show the most rearrangements compared to the other *Chloropicon* taxa. Surprisingly, it was found that only three reversals separate the *Chloropicon maureeniae* and *Chloroparvula* mitogenomes. With 30 to 32 reversals, the mitogenome of *Chloropicon* sp. RCC4434 differs as extensively from the other mitogenomes of the *Chloropicon* lineage as from the *Chloroparvula* mitogenomes.

As reported above for the plastomes, the mitogenome sequences of closely related taxa could be aligned entirely or almost entirely using LAST and again, *Chloroparvula* sp. RCC696 and RCC4572 displayed the lowest density of SNPs, followed by the *Chloropicon roscoffensis* strains (fig. 8 and supplementary table S2, Supplementary Material online).

## Discussion

### Organelle phylogenomic analyses provide a robust phylogeny of the Chloropicophyceae

Given their ecological importance in open oceanic waters (Lopes dos Santos, et al. 2017a), the tiny green algae from the newly erected prasinophyte class Chloropicophyceae are bound to be the targets of future investigations aimed at better understanding their genetic divergence, biology, habitats, and role in ecosystems. In this context, knowledge about the precise relationships among the various species or clades (A1-A7 and B1-B3) in each of the two genera of this class is essential, as it provides the evolutionary framework required to establish the timing of changes that occurred during species diversification. The plastome- and mitogenome-based phylogenomic trees we report in the present study resolved with robustness the relationships among the *Chloropicon* and *Chloroparvula* species (fig. 1). These trees exhibit exactly the same topology but differ from the weakly supported tree of the Chloropicophyceae inferred from concatenated nuclear 18S and plastid 16S rRNA genes (Lopes dos Santos, et al. 2017b) with regards to the affinities between *Chloropicon laureae* (A5) and *Chloropicon mariensis* (A1) as well as the relationships among the three closely related *Chloroparvula* sp. taxa originally assigned to the B2 and B3 lineages. It should be considered that the *Chloropicon* species from the A1 and A5 lineages share a common ancestor, a conclusion also congruent with the robust relationships inferred for *Chloropicon* species from the predicted amino acid sequences of transcripts corresponding to 127 nuclear genes (Lopes dos Santos, et al. 2017b); however, it remains unclear why the results reported here differ from the 18S+16S rRNA tree in showing a closer affinity between the B2 RCC696 and B3 RCC4572 strains than between the B2 RCC696 and RCC699 strains (figs. 1 and 8). Importantly, the plastome-based phylogenomic analysis including representatives of other Chloroplastida lineages revealed that the *Chloropicon* and *Chloroparvula* genera are separated by a very long evolutionary distance and that each chloropicophycean genus shows a substantially greater level of sequence divergence compared to the mamiellalean genera *Ostreococcus* and *Micromonas* (fig. 2). The same global phylogenomic tree confirmed that the Chloropicophyceae are sister to all core chlorophytes; however, the prasinophyte lineage that diverged just before the Chloropicophyceae could not be identified, as the relative positions of the Picocystophyceae and Pycnococcaceae were not clearly resolved.

### Organelle genomes sustained extensive changes during the evolution of the Chloropicophyceae

Consistent with the high level of sequence divergence separating the *Chloropicon* and *Chloroparvula* genera, considerable architectural alterations were identified in the plastomes and mitogenomes assembled in the present study, and their mapping on the inferred phylogenomic trees provided significant insights into the evolutionary histories of organelle genomes in the Chloropicophyceae (fig. 9). So far, only a few studies tracked in a comprehensive manner structural genomic changes in the chloroplasts and mitochondria of green algae from the same lineage (Smith and Keeling 2015; Turmel, et al. 2016). Although the evolutionary pattern observed for one or the other organelle genome is typically variable across green algal lineages, shared patterns have been observed for the plastome and mitogenome of certain lineages of core chlorophytes, in particular with regards to the co-occurrence of important variations in A+T content and in the proportion of noncoding sequences (Smith and Keeling 2015). In contrast, it is well documented that the plastome and mitogenome of land plants, in particular angiosperms, have their own characteristic pattern, with the mitogenome evolving much more slowly at the sequence level than the plastome and also rearranging more frequently (Mower, et al. 2012; Mower and Vickrey 2018).

**Fig. 9.**
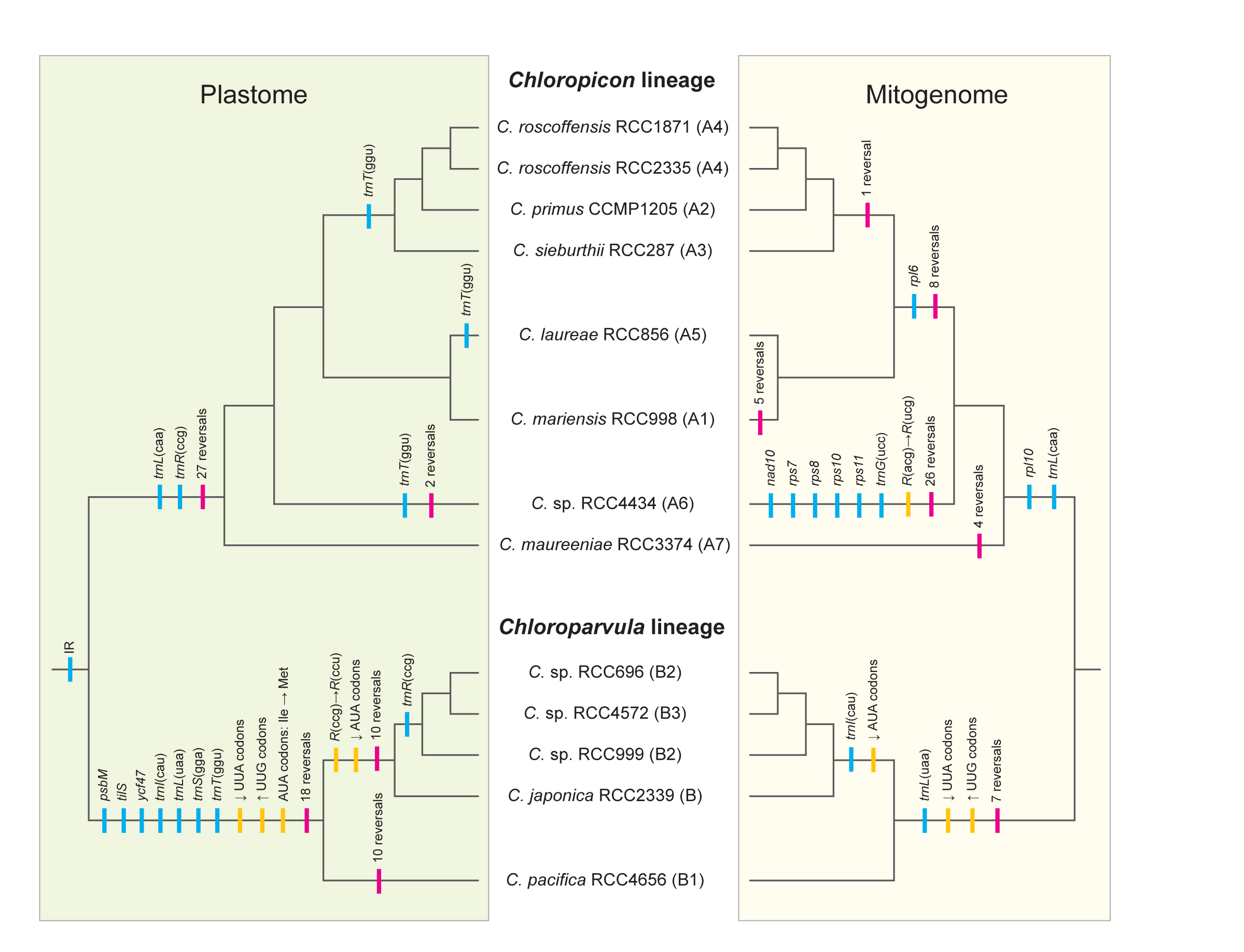
Predicted evolutionary scenarios for the plastome and mitogenome of the Chloropicophyceae. Gene losses and gene rearrangements are denoted by blue and red vertical lines, respectively. Mutations of tRNA anticodons, changes in codon frequencies, and reassignment of the AUA codon are indicated by yellow lines.

The plastomes and mitogenomes of the Chloropicophyceae experienced similar types of changes, but the tempos of occurrence of these changes differ in the two organelles (fig. 9). In the plastid, numerous gene losses and extensive gene reshuffling accompanied the split of the *Chloropicon* and *Chloroparvula* lineages; however, the plastome changed at a much slower pace during species diversification in each genus. In contrast, the mitogenome sustained more gradual changes during the evolution of the Chloropicophyceae and less extensive rearrangements during the emergence of the *Chloropicon* and *Chloroparvula* lineages. Differential losses of three mitochondrial genes [*rpl10*, *trnL*(uaa) and *trnL*(caa)] coincided with the divergence of the *Chloropicon* and *Chloroparvula* lineages, and further losses of *rpl6* and *trnI*(cau) took place following the branching off of the *Chloropicon maureeniae* and *Chloroparvula pacifica* lineages. Remarkably, both the plastome and mitogenome lost three tRNA genes with the same decoding functions [*trnI*(cau), *trnL*(uaa) and *trnL*(caa)], leading to important changes in codon usage (see next section).

Prior to our study, the plastome of *Chloropicon primus* was the only organelle genome of the Chloropicophyceae that was available for comparison with its homologs in other prasinophyte lineages (Lemieux, et al. 2014). Like this plastome, the newly sequenced plastomes of *Chloroparvula* are extremely divergent from all those previously sampled from prasinophytes, revealing no phylogenetic link with a specific prasinophyte lineage as well as no clue as to what the plastome architecture looked like during the early evolution of the Chloropicophyceae. Considering that the *Picocystis salinarum* plastome (Lemieux, et al. 2014) is substantially more gene-rich than the *Chloroparvula* and *Chloropicon* plastomes and has retained a rRNA operon-containing IR, it is likely that ancestral traits have massively disappeared soon after the emergence of the Chloropicophyceae. But the situation appears different for the mitogenomes of the Chloropicophyceae, as their gene repertoires are similar to those found in *Picocystis* (table 1, supplementary fig. S6, Supplementary Material Online), the Mamiellophyceae (Robbens, et al. 2007) and Nephroselmidophyceae (Turmel, et al. 1999a). As observed for the plastome, however, they are lacking an IR containing the rRNA genes, a feature that is probably an ancestral trait given its presence in *Picocystis*, the Mamiellophyceae and *Prasinoderma coloniale* (Pombert, et al. 2013; Robbens, et al. 2007).

Following the divergence of the *Chloropicon* and *Chloroparvula* lineages, the plastome maintained a nearly identical size and gene density in each genus, whereas the mitogenome fluctuated at these levels (table 1). It is not clear what are the causes of these differences: we cannot exclude the possibility that there is an elevated pressure to keep the plastome more reduced and gene-dense than the mitogenome, but there might also exist differences related to the type and rate of errors associated with the mechanisms ensuring DNA integrity in the two organelles. As proposed for land plants (Christensen 2014), DNA repair may be more prone to errors in the mitogenome than the plastome in the Chloropicophyceae, thus possibly accounting for the lower gene density found in the mitogenome. Considering that gene rearrangements are the products of illegitimate recombination events involving different intergenic regions, organelle genomes with longer intergenic regions, which is the case here for the mitogenome, are expected to have more opportunity to recombine. Consistent with this idea, the mitogenomes of representatives from several *Chloropicon* clades exhibited substantial gene reversals (fig. 9). The mitogenome of *Chloropicon* sp. RCC4434, which is the longest in the *Chloropicon* lineage, was the most rearranged relative to its counterparts; however, the *Chloroparvula* mitogenomes, including the largest identified in the Chloropicophyceae (*Chloroparvula pacifica* mitogenome) were all colinear. The seemingly higher rate of mitogenome recombination in the *Chloropicon* genus may be due to differences in the efficiency of the nuclear-encoded enzymes catalyzing recombination or to the presence of more favourable sequences serving as DNA substrates for these enzymes.

Recent studies comparing substitution rates in organelle genomes have shown that the mitogenome is subjected to a higher mutation rate than the plastome in a number of algal lineages (Smith 2015). In green algae, this trend has been noted for *Ostreococcus* but not for *Chlamydomonas* (Chlorophyceae) and the streptophyte *Mesostigma*. Our analyses of chloropicophycean plastomes and mitogenomes also revealed a general trend toward a higher mutation rate in the mitogenome, even though the relative substitution rate between the two organelle genomes varied depending on the *Chloroparvula* isolates examined (fig. 8).

### Adaptation of codon usage in *Chloroparvula* plastomes and mitogenomes, with reassignment of the AUA codon to methionine in the plastome

Losses of genes from organelle genomes may have diverse consequences depending on the function of the encoded products. Before being completely extinguished, organelle genes essential for cell survival are generally transferred to the nucleus where they are endowed with the presequence necessary for organelle targeting of their encoded products or alternatively, they are replaced by existing nuclear genes of similar function whose products are retargeted to the organelle. Such events of gene transfer to the nucleus are predicted for the photosynthetic gene *psbM* and the ribosomal-protein genes that were found to be missing in some strains of the Chloropicophyceae (figs. 3A and 3B), but firm evidence for their occurrence will await characterization of nuclear genomes from these strains. While tRNA genes with dispensable or redundant function can disappear from the cell without any further impact, loss of genes encoding tRNAs needed for reading specific codons leads to adaptation of codon usage (figs. 3C-3F). To our knowledge, our study provides the first evidence for codon adaptation resulting from losses of *trnI*(cau) and *trnL*(uaa) in algae. Remarkably, these two genes, which are almost invariably present in Chloroplastida organelle genomes, suffered convergent loss from both the plastome and mitogenome in the *Chloroparvula* lineage and these events were intimately associated with the disappearance of the original AUA isoleucine and UUA leucine codons recognized by their encoded tRNAs. These codons vanished through mutations mainly to AUU isoleucine and UUG leucine codons, respectively (supplementary table S1, Supplementary Material Online).

The status of the unassigned AUA codons was temporary following the loss of *trnI*(cau) and *tilS* in plastomes, as new AUA codons were created by mutations of AUG codons in the *Chloroparvula* lineage and translated to methionine residues following the emergence of a modified tRNA^Met^(CAU). The frequent occurrence of these codons at universally or almost universally conserved sites exhibiting AUG codons in plastomes of phylogenetically diverse green algae together with the complete lack of AUA codons at highly conserved codon sites decoded as isoleucine provide compelling evidence not only for the evolution of a variant genetic code in which both AUA and AUG are read as methionine (fig. 5 and supplementary fig. S4, Supplementary Material Online) but also for the mechanism proposed above, which is also known as the codon-capture hypothesis (Osawa, et al. 1989; Watanabe and Yokobori 2011). This variant code is used in the mitochondria of most metazoans and some yeasts (*Sacharomyces* and *Torulopsis*) and the codon capture hypothesis was also favored to explain the reassignment of AUA to methionine in these mitochondria (Osawa, et al. 1989). To our knowledge, this is the first time that this noncanonical genetic code is reported in plastids. In green algal plastomes, a variant genetic code has been previously documented only for the chlorophyte *Jenufa minuta* (Sphaeropleales, Chlorophyceae), where UGA codons, which are normally stop codons, specify tryptophan (Turmel and Lemieux 2018).

Efficient decoding of the reintroduced AUA codons as methionine required alterations of an existing tRNA^Met^(CAU) in metazoan mitochondria (Watanabe and Yokobori 2011). Indeed, it has been shown that a modified nucleotide in the anticodon loop at the wobble position 34 (5-formylcytidine) or at position 37 (N^6^-threonylcarbamyladenosine) enables the tRNA^Met^(CAU) to pair with both AUA and AUG codons. Although no information is currently available about the presence of modified nucleotides at these positions in plastid tRNAs^Met^(CAU), alignments of the *Chloroparvula* tRNA gene sequences with the corresponding sequences of *Chloropicon* and other green algae unveiled specific substitutions in the former sequences that map to three of the five base-pairings forming the stem of the anticodon arm (fig. 6). The changes in the C29:G41 and U31:G29 pairs are especially noteworthy considering that these base-pairs are very conserved among chlorophyte tRNAs^Met^(CAU). Perhaps these mutations may influence the anticodon-codon interaction and allow the wobble position of the anticodon to recognize both purine nucleotides at the third codon position. Obviously, to decipher the mechanism that mediated the reassignment of the AUA codon in the Chloropicophyceae, it will be important to investigate whether the *Chloroparvula* and *Chloropicon* tRNAs^Met^(CAU) differ with respect to modified nucleotides in the anticodon loop.

## Acknowledgements

This work was supported by the Natural Sciences and Engineering Research Council of Canada (http://www.nserc-crsng.gc.ca/index_eng.asp) (Grant RGPIN-2017-04506 to CL). We thank Daniel Vaulot and Ehsan Kayal for helpful discussions.

## References

Alkatib S, et al. 2012. The contributions of wobbling and superwobbling to the reading of the genetic code. PLoS Genet 8: e1003076.

Bankevich A, et al. 2012. SPAdes: a new genome assembly algorithm and its applications to single-cell sequencing. J Comput Biol 19: 455–477.

Bourque G, Pevzner PA 2002. Genome-scale evolution: reconstructing gene orders in the ancestral species. Genome Res 12: 26–36.

Cahoon AB, Nauss JA, Stanley CD, Qureshi A 2017. Deep transcriptome sequencing of two green algae, Chara vulgaris and Chlamydomonas reinhardtii, provides no evidence of organellar RNA editing. Genes 8.

Capella-Gutierrez S, Silla-Martinez JM, Gabaldon T 2009. trimAl: a tool for automated alignment trimming in large-scale phylogenetic analyses. Bioinformatics 25: 1972–1973.

Castresana J 2000. Selection of conserved blocks from multiple alignments for their use in phylogenetic analysis. Mol Biol Evol 17: 540–552.

Chen S, et al. 2017. AfterQC: automatic filtering, trimming, error removing and quality control for fastq data. BMC Bioinformatics 18: 80.

Christensen AC 2014. Genes and junk in plant mitochondria—repair mechanisms and selection. Genome Biol Evol 6: 1448–1453.

Darling AE, Mau B, Perna NT 2010. progressiveMauve: multiple genome alignment with gene gain, loss and rearrangement. PLoS One 5: e11147.

Edgar RC 2004. MUSCLE: multiple sequence alignment with high accuracy and high throughput. Nucleic Acids Res 32: 1792–1797.

Frith MC, Hamada M, Horton P 2010. Parameters for accurate genome alignment. BMC Bioinformatics 11: 80.

Gitzendanner MA, Soltis PS, Yi T-S, Li D-Z, Soltis DE 2018. Plastome Phylogenetics: 30 Years of Inferences Into Plant Evolution. Adv Bot Res 85: 293–313.

Grimsley N, Yau S, Piganeau G, Moreau H. 2015. Typical features of genomes in the Mamiellophyceae. In: Ohtsuka S, Suzaki T, Horiguchi T, Suzuki N, Not F, editors. Marine Protists: Diversity and Dynamics. Tokyo: Springer Japan. p. 107–127.

Guillou L, et al. 2004. Diversity of picoplanktonic prasinophytes assessed by direct nuclear SSU rDNA sequencing of environmental samples and novel isolates retrieved from oceanic and coastal marine ecosystems. Protist 155: 193–214.

Hoang DT, Chernomor O, von Haeseler A, Minh BQ, Vinh LS 2018. UFBoot2: Improving the Ultrafast Bootstrap Approximation. Mol Biol Evol 35: 518–522.

Hrda S, Hroudova M, Vlcek C, Hampl V 2017. Mitochondrial genome of prasinophyte alga Pyramimonas parkeae. J Eukaryot Microbiol 64: 360–369.

Hu F, Lin Y, Tang J 2014. MLGO: phylogeny reconstruction and ancestral inference from geneorder data. BMC Bioinformatics 15: 354.

Kikuchi S, et al. 2013. Uncovering the protein translocon at the chloroplast inner envelope membrane. Science 339: 571–574.

Lartillot N, Lepage T, Blanquart S 2009. PhyloBayes 3: a Bayesian software package for phylogenetic reconstruction and molecular dating. Bioinformatics 25: 2286–2288.

Leliaert F, et al. 2016. Chloroplast phylogenomic analyses reveal the deepest-branching lineage of the Chlorophyta, Palmophyllophyceae class. nov. Sci Rep 6: 25367.

Lemieux C, Otis C, Turmel M 2014. Six newly sequenced chloroplast genomes from prasinophyte green algae provide insights into the relationships among prasinophyte lineages and the diversity of streamlined genome architecture in picoplanktonic species. BMC Genomics 15: 857.

Lohse M, Drechsel O, Kahlau S, Bock R 2013. OrganellarGenomeDRAW--a suite of tools for generating physical maps of plastid and mitochondrial genomes and visualizing expression data sets. Nucleic Acids Res 41: W575–581.

Lopes dos Santos A, Gourvil P, Rodriguez F, Garrido JL, Vaulot D 2016. Photosynthetic pigments of oceanic Chlorophyta belonging to prasinophytes clade VII. J Phycol 52: 148–155.

Lopes dos Santos A, et al. 2017a. Diversity and oceanic distribution of prasinophytes clade VII, the dominant group of green algae in oceanic waters. ISME J 11: 512–528.

Lopes dos Santos A, et al. 2017b. Chloropicophyceae, a new class of picophytoplanktonic prasinophytes. Sci Rep 7: 14019.

Lowe TM, Eddy SR 1997. tRNAscan-SE: a program for improved detection of transfer RNA genes in genomic sequence. Nucleic Acids Res 25: 955–964.

Maddison WP, Maddison DR 2018. Mesquite: a modular system for evolutionary analysis. Version 3.51. http://mesquiteproject.org.

Moreau H, et al. 2012. Gene functionalities and genome structure in Bathycoccus prasinos reflect cellular specializations at the base of the green lineage. Genome Biol 13: R74.

Mower JP, Sloan DB, Alverson AJ 2012. Plant mitochondrial genome diversity: the genomics revolution. In: Wendel JH, editor. Plant genome diversity volume 1: Plant genomes, their residents, and their evolutionary dynamics. New York: Springer. p. 123–124.

Mower JP, Vickrey TL 2018. Structural diversity among plastid genomes of land plants. Adv Bot Res 85: 263–292.

Nguyen LT, Schmidt HA, von Haeseler A, Minh BQ 2015. IQ-TREE: a fast and effective stochastic algorithm for estimating maximum-likelihood phylogenies. Mol Biol Evol 32: 268–274.

Osawa S, Ohama T, Jukes TH, Watanabe K, Yokoyama S 1989. Evolution of the mitochondrial genetic code. II. Reassignment of codon AUA from isoleucine to methionine. J Mol Evol 29: 373–380.

Pombert JF, Otis C, Turmel M, Lemieux C 2013. The mitochondrial genome of the prasinophyte Prasinoderma coloniale reveals two trans-spliced group I introns in the large subunit rRNA gene. PLoS One 8: e84325.

Potter D, Lajeunesse TC, Saunders GW, Anderson RA 1997. Convergent evolution masks extensive biodiversity among marine coccoid picoplankton. Biodiversity & Conservation 6: 99–107.

Rice P, Longden I, Bleasby A 2000. EMBOSS: The European molecular biology open software suite. Trends Genet 16: 276–277.

Rii YM, et al. 2016. Diversity and productivity of photosynthetic picoeukaryotes in biogeochemically distinct regions of the South East Pacific Ocean. Limnology and Oceanography 61: 806–824.

Robbens S, et al. 2007. The complete chloroplast and mitochondrial DNA sequence of Ostreococcus tauri: organelle genomes of the smallest eukaryote are examples of compaction. Mol Biol Evol 24: 956–968.

Satjarak A, Burns JA, Kim E, Graham LE 2017. Complete mitochondrial genomes of prasinophyte algae Pyramimonas parkeae and Cymbomonas tetramitiformis. J Phycol 53: 601–615.

Satjarak A, Graham LE 2017. Comparative DNA sequence analyses of Pyramimonas parkeae (Prasinophyceae) chloroplast genomes. J Phycol 53: 415–424.

Smith DR 2015. Mutation rates in plastid genomes: they are lower than you might think. Genome Biol Evol 7: 1227–1234.

Smith DR, Keeling PJ 2015. Mitochondrial and plastid genome architecture: Reoccurring themes, but significant differences at the extremes. Proc Natl Acad Sci U S A 112: 10177–10184.

Smith SA, Dunn CW 2008. Phyutility: a phyloinformatics tool for trees, alignments and molecular data. Bioinformatics 24: 715–716.

Soma A, et al. 2003. An RNA-modifying enzyme that governs both the codon and amino acid specificities of isoleucine tRNA. Mol Cell 12: 689–698.

Stamatakis A 2014. RAxML version 8: a tool for phylogenetic analysis and post-analysis of large phylogenies. Bioinformatics 30: 1312–1313.

Sym SD. 2015. Basal lineages of green algae – Their diversity and phylogeny. In: Ohtsuka S, Suzaki T, Horiguchi T, Suzuki N, Not F, editors. Marine Protists: Diversity and Dynamics. Tokyo: Springer Japan. p. 89–105.

Tesler G 2002. GRIMM: genome rearrangements web server. Bioinformatics 18: 492–493.

Tragin M, Vaulot D 2018. Green microalgae in marine coastal waters: The Ocean Sampling Day (OSD) dataset. Sci Rep 8: 14020.

Turmel M, Gagnon MC, O’Kelly CJ, Otis C, Lemieux C 2009. The chloroplast genomes of the green algae Pyramimonas, Monomastix, and Pycnococcus shed new light on the evolutionary history of prasinophytes and the origin of the secondary chloroplasts of euglenids. Mol Biol Evol 26: 631–648.

Turmel M, Lemieux C 2018. Evolution of the plastid genome in green algae. Adv Bot Res 85: 157–193.

Turmel M, et al. 1999a. The complete mitochondrial DNA sequences of *Nephroselmis olivacea and Pedinomonas minor*. Two radically different evolutionary patterns within green algae. Plant Cell 11: 1717–1730.

Turmel M, Otis C, Lemieux C 1999b. The complete chloroplast DNA sequence of the green alga Nephroselmis olivacea: insights into the architecture of ancestral chloroplast genomes. Proc Natl Acad Sci U S A 96: 10248–10253.

Turmel M, Otis C, Lemieux C 2010. A deviant genetic code in the reduced mitochondrial genome of the picoplanktonic green alga Pycnococcus provasolii. J Mol Evol 70: 203–214.

Turmel M, Otis C, Lemieux C 2017. Divergent copies of the large inverted repeat in the chloroplast genomes of ulvophycean green algae. Sci Rep 7: 994.

Turmel M, Otis C, Lemieux C 2016. Mitochondrion-to-Chloroplast DNA transfers and intragenomic proliferation of chloroplast group II introns in Gloeotilopsis green algae (Ulotrichales, Ulvophyceae). Genome Biol Evol 8: 2789–2805.

Watanabe K, Yokobori S 2011. tRNA modification and genetic code variations in animal mitochondria. J Nucleic Acids 2011: 623095.

Waterhouse AM, Procter JB, Martin DM, Clamp M, Barton GJ 2009. Jalview Version 2-a multiple sequence alignment editor and analysis workbench. Bioinformatics 25: 1189–1191.

Worden AZ, et al. 2009. Green evolution and dynamic adaptations revealed by genomes of the marine picoeukaryotes Micromonas. Science 324: 268–272.

Yu M, et al. 2018. Evolution of the Plastid Genomes in Diatoms. Adv Bot Res 85: 129–155.

