## Supplementary_figures_S1-S6 for "Tracing the evolution of the plastome and mitogenome in the Chloropicophyceae uncovered convergent tRNA gene losses and a variant plastid genetic code"

**Fig. S1.**—Gene maps of all plastomes compared in this study, except that previously reported for *Chloropicon primus* CCMP1205 (Lemieux, et al. 2014). Filled boxes represent genes, with colors denoting gene categories as indicated in the legend. Genes on the outside of the map are transcribed counterclockwise; those on the inside are transcribed clockwise. The inner ring shows variations in G+C content, with the circle inside the G+C content graph marking the 50% threshold (dark gray, G+C; light gray, A+T). GenBank accession numbers of all plastomes are indicated in Table 1.

**Fig. S2.**—Gene maps of all mitogenomes compared in this study, except that of *Chloropicon primus* CCMP1205 which is shown in supplementary fig. S6, Supplementary Material online. Filled boxes represent genes, with colors denoting gene categories as indicated in the legend. Genes on the outside of the map are transcribed counterclockwise; those on the inside are transcribed clockwise. The inner ring shows variations in G+C content, with the circle inside the G+C content graph marking the 50% threshold (dark gray, G+C; light gray, A+T). GenBank accession numbers of all mitogenomes are indicated in Table 1.

**Fig. S3.**—Multiple sequence alignments of Ycf1 proteins encoded by the plastomes of chloropicophycean taxa and *Picocystis salinarum*.

**Fig. S4.**—*chlB* gene alignment showing that no AUA codons (shaded in black) in *Chloroparvula* plastomes correspond to universally or almost universally conserved isoleucine residues (red columns) in the encoded protein. Instead, all AUA codons falling within universally or almost universally conserved amino acid residues correspond to methionine AUG codons (blue columns).

**Fig. S5.**—Estimated numbers of reversals in pairwise comparisons of chloropicophycean plastomes and mitogenomes using GRIMM.

**Fig. S6.**—Gene maps of *Picocystis salinarum* and *Chloropicon primus* mitogenomes and comparison of gene content between these genomes. Only the genes missing in one or the other genome are shown, with gene presence denoted by a blue box. The following 59 genes are shared by both mitogenomes: *atp1,4,6,8,9, cob, cox1,2,3, mttB, nad1,2,3,4,4L,5,6,7,9,10, rpl5,14,16, rps2,3,4,7,8,10,11,12,13,14,19, rnl, rns, trnA*(ugc), *C*(gca), *D*(guc), *E*(uuc), *F*(gaa), *G*(gcc), *G*(ucc), *H*(gug), *I*(gau), *K*(uuu), L(uag), *Me*(cau), *Mf*(cau), *N*(guu), *P*(ugg), *Q*(uug), *R*(acg), *R*(ucu), *S*(gcu), *S*(uga), *V*(uac), *W*(cca), *Y*(gua). GenBank accession numbers of the two compared mitogenomes are indicated in Table 1.

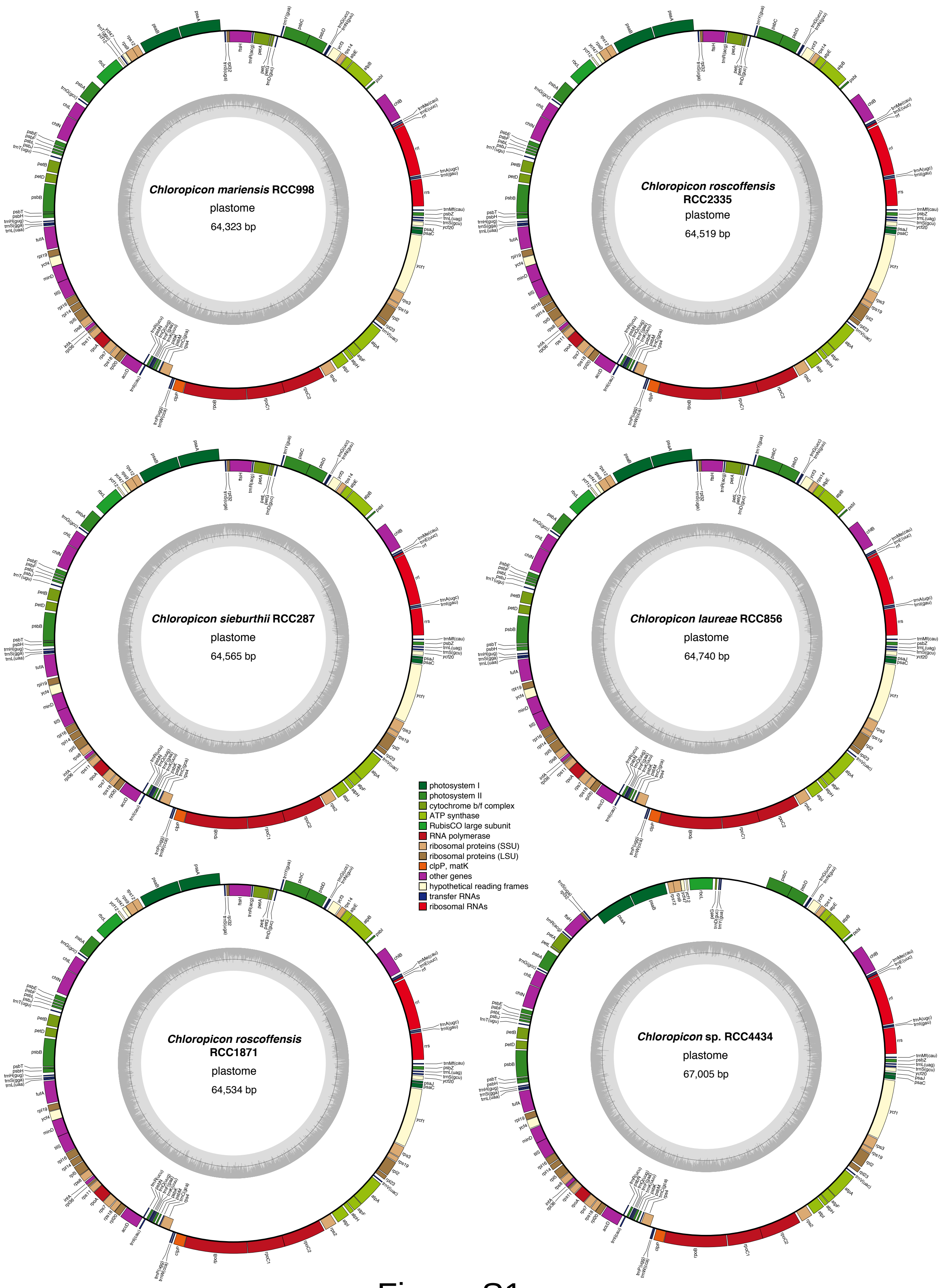
*Chloropicon mariensis* RCC998
plastome
64,323 bp
*Chloropicon roscoffensis* RCC2335
plastome
64,519 bp
*Chloropicon sieburthii* RCC287
plastome
64,565 bp
*Chloropicon laureae* RCC856
plastome
64,740 bp
*Chloropicon roscoffensis* RCC1871
plastome
64,534 bp
*Chloropicon* sp. RCC4434
plastome
67,005 bp
photosystem I
photosystem II
cytochrome b/f complex
ATP synthase
RubisCO large subunit
RNA polymerase
ribosomal proteins (SSU)
ribosomal proteins (LSU)
clpP, matK
other genes
hypothetical reading frames
transfer RNAs
ribosomal RNAs

Figure S1

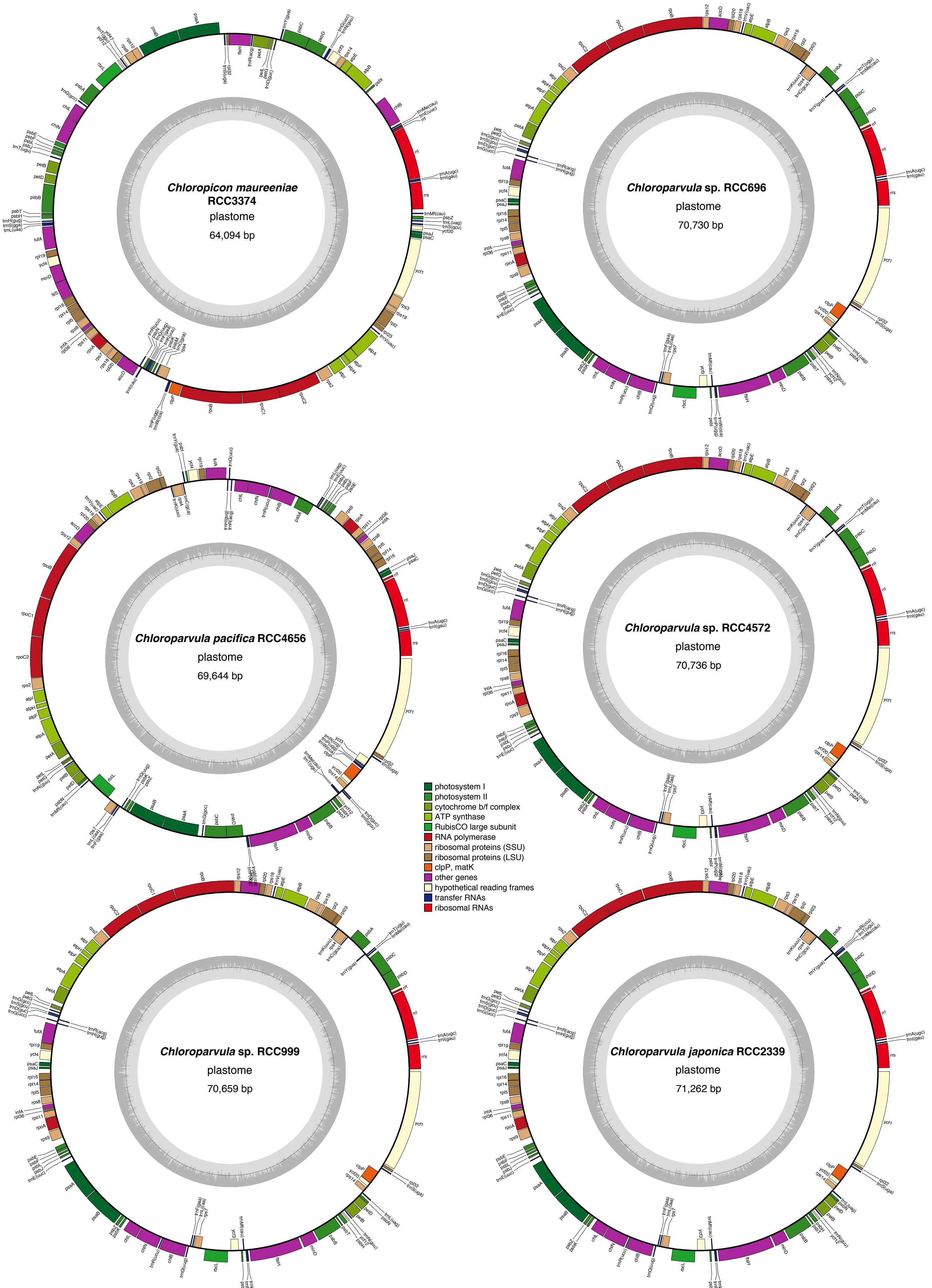
Chloropicon maureeniae RCC3374
plastome
64,094 bp
Chloroparvula sp. RCC696
plastome
70,730 bp
Chloroparvula pacifica RCC4656
plastome
69,644 bp
Chloroparvula sp. RCC4572
plastome
70,736 bp
Chloroparvula sp. RCC999
plastome
70,659 bp
Chloroparvula japonica RCC2339
plastome
71,262 bp
photosystem I
photosystem II
cytochrome b/f complex
ATP synthase
RubisCO large subunit
RNA polymerase
ribosomal proteins (SSU)
ribosomal proteins (LSU)
clpP, matK
other genes
hypothetical reading frames
transfer RNAs
ribosomal RNAs

Figure S1

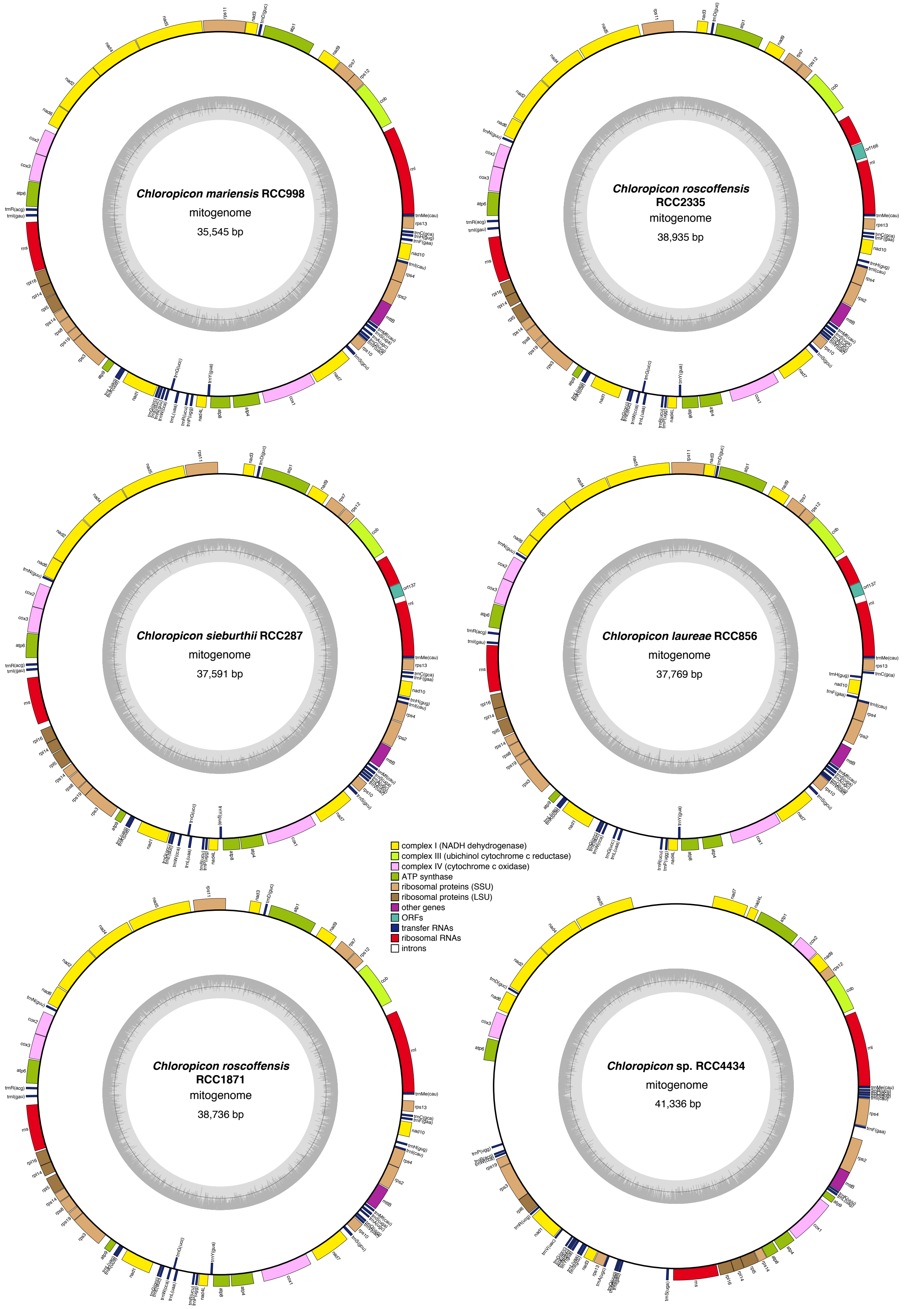

Chloropicon mariensis RCC998
mitogenome
35,545 bp
Chloropicon roscoffensis
RCC2335
mitogenome
38,935 bp
Chloropicon sieburthii RCC287
mitogenome
37,591 bp
Chloropicon laureae RCC856
mitogenome
37,769 bp
Chloropicon roscoffensis
RCC1871
mitogenome
38,736 bp
Chloropicon sp. RCC4434
mitogenome
41,336 bp
complex I (NADH dehydrogenase)
complex III (ubichinol cytochrome c reductase)
complex IV (cytochrome c oxidase)
ATP synthase
ribosomal proteins (SSU)
ribosomal proteins (LSU)
other genes
ORFs
transfer RNAs
ribosomal RNAs
introns


Figure S2

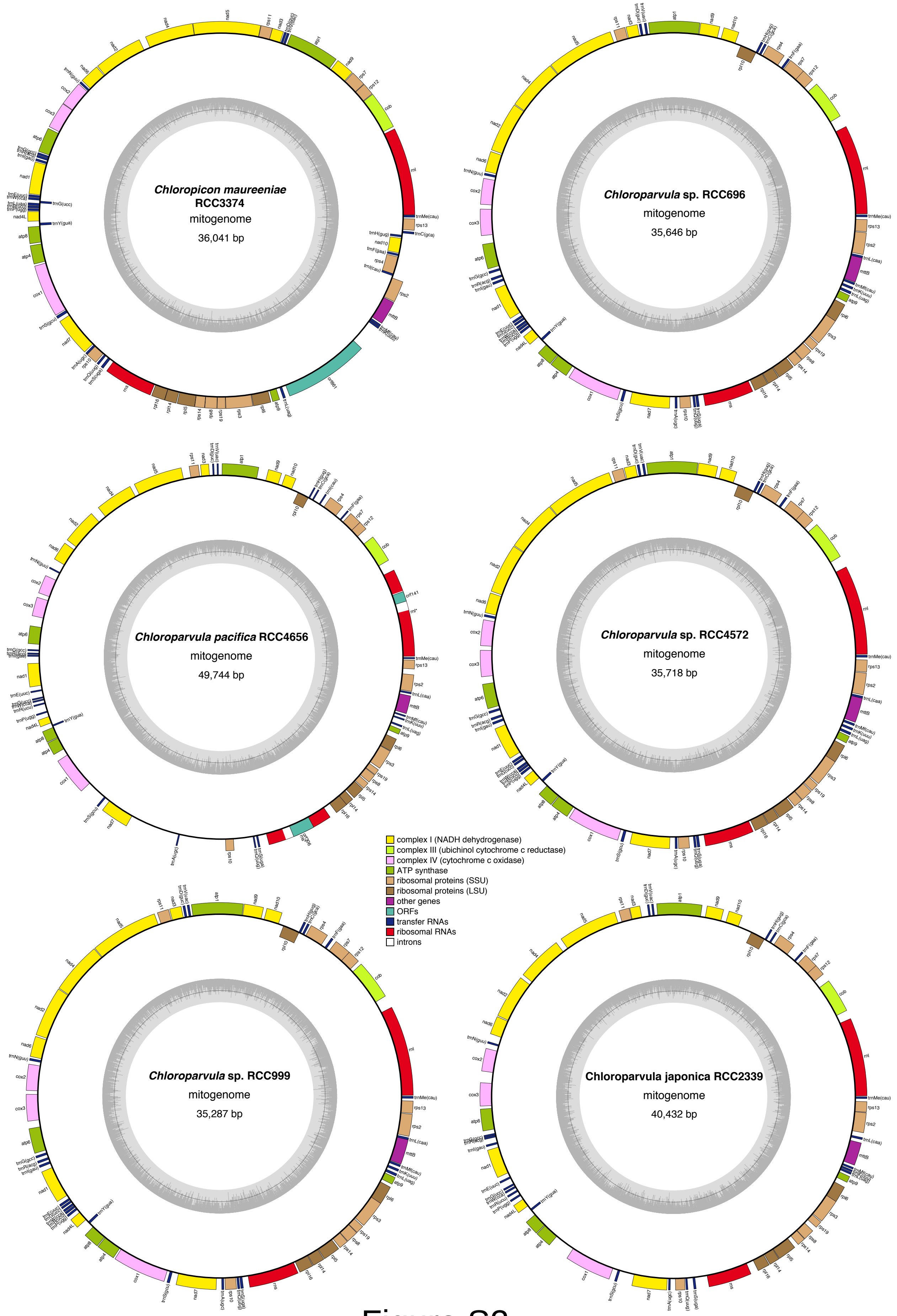
Chloropicon maureeniae RCC3374
mitogenome
36,041 bp
Chloroparvula sp. RCC696
mitogenome
35,646 bp
Chloroparvula pacifica RCC4656
mitogenome
49,744 bp
Chloroparvula sp. RCC4572
mitogenome
35,718 bp
Chloroparvula sp. RCC999
mitogenome
35,287 bp
Chloroparvula japonica RCC2339
mitogenome
40,432 bp
complex I (NADH dehydrogenase)
complex III (ubichinol cytochrome c reductase)
complex IV (cytochrome c oxidase)
ATP synthase
ribosomal proteins (SSU)
ribosomal proteins (LSU)
other genes
ORFs
transfer RNAs
ribosomal RNAs
introns

Figure S2

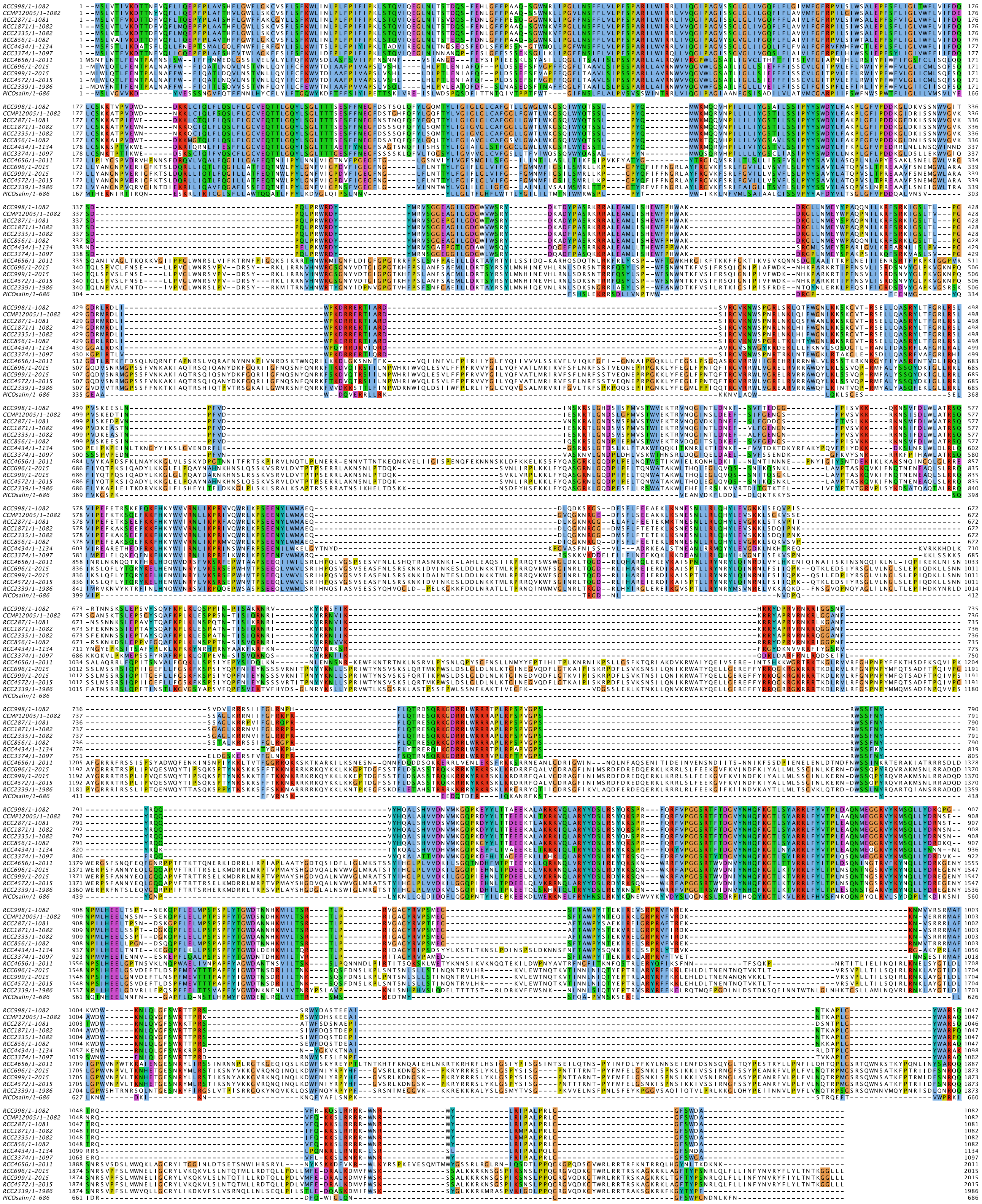

Figure S3

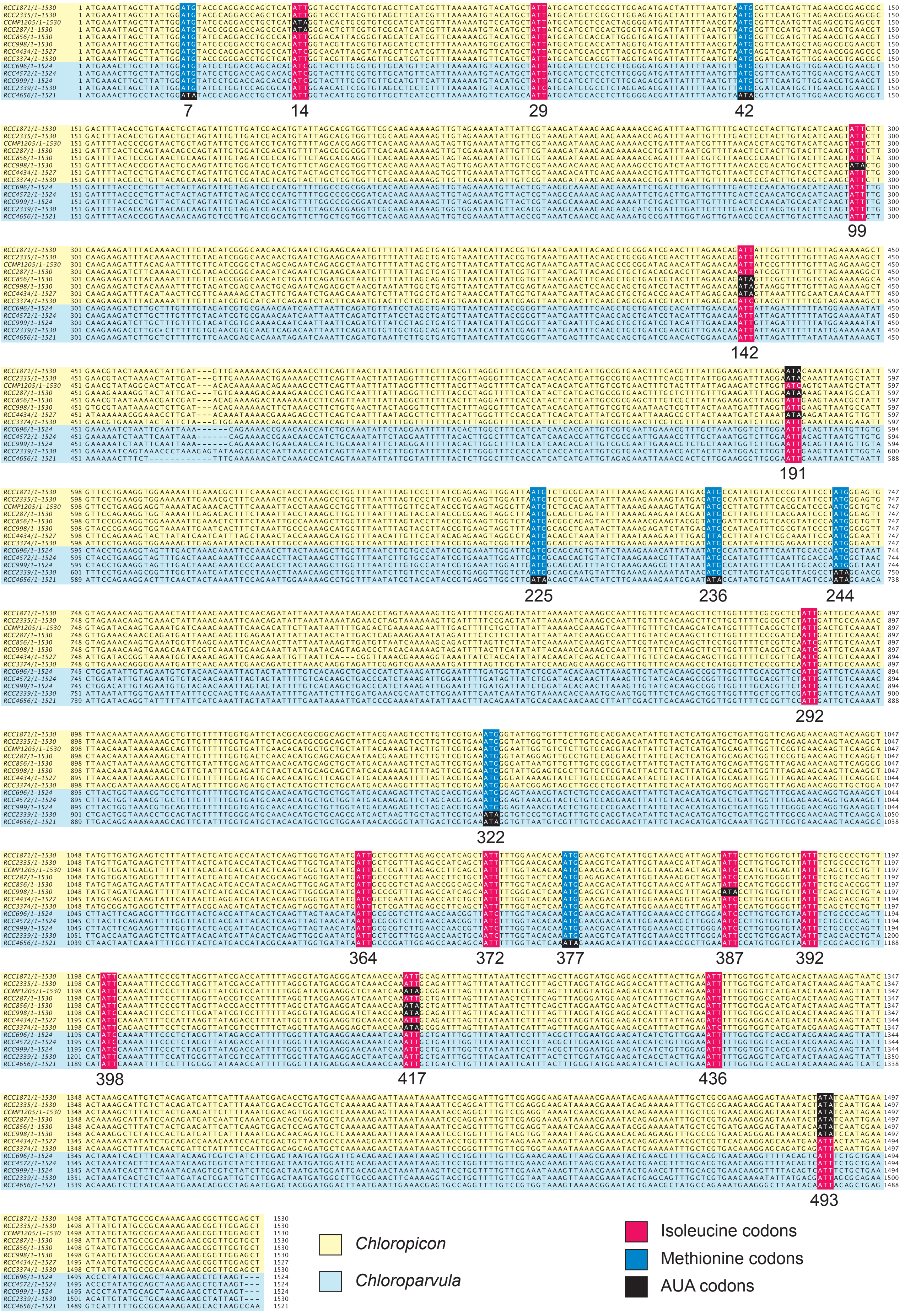
7
14
29
42
99
142
191
225
236
244
292
322
364
372
377
387
392
398
417
436
493
Chloropicon
Chloroparvula
Isoleucine codons
Methionine codons
AUA codons

Figure S4

### A. Plastome

13 genomes, 95 genes — *Chloropicon* — *Chloroparvula*

| | RCC998 | CCMP1205 | RCC287 | RCC1871 | RCC2335 | RCC856 | RCC3374 | RCC4434 | RCC999 | RCC4572 | RCC696 | RCC2339 | RCC4656 |
|---|---|---|---|---|---|---|---|---|---|---|---|---|---|
| RCC998 | 0 | 0 | 0 | 0 | 0 | 0 | 0 | 2 | 53 | 53 | 53 | 53 | 53 |
| CCMP1205 | 0 | 0 | 0 | 0 | 0 | 0 | 0 | 2 | 53 | 53 | 53 | 53 | 53 |
| RCC287 | 0 | 0 | 0 | 0 | 0 | 0 | 0 | 2 | 53 | 53 | 53 | 53 | 53 |
| RCC1871 | 0 | 0 | 0 | 0 | 0 | 0 | 0 | 2 | 53 | 53 | 53 | 53 | 53 |
| RCC2335 | 0 | 0 | 0 | 0 | 0 | 0 | 0 | 2 | 53 | 53 | 53 | 53 | 53 |
| RCC856 | 0 | 0 | 0 | 0 | 0 | 0 | 0 | 2 | 53 | 53 | 53 | 53 | 53 |
| RCC3374 | 0 | 0 | 0 | 0 | 0 | 0 | 0 | 2 | 53 | 53 | 53 | 53 | 53 |
| RCC4434 | 2 | 2 | 2 | 2 | 2 | 2 | 2 | 0 | 53 | 53 | 53 | 53 | 53 |
| RCC999 | 53 | 53 | 53 | 53 | 53 | 53 | 53 | 53 | 0 | 0 | 0 | 0 | 19 |
| RCC4572 | 53 | 53 | 53 | 53 | 53 | 53 | 53 | 53 | 0 | 0 | 0 | 0 | 19 |
| RCC696 | 53 | 53 | 53 | 53 | 53 | 53 | 53 | 53 | 0 | 0 | 0 | 0 | 19 |
| RCC2339 | 53 | 53 | 53 | 53 | 53 | 53 | 53 | 53 | 0 | 0 | 0 | 0 | 19 |
| RCC4656 | 53 | 53 | 53 | 53 | 53 | 53 | 53 | 53 | 19 | 19 | 19 | 19 | 0 |

### B. Mitogenome

13 genomes, 53 genes — *Chloropicon* — *Chloroparvula*

| | RCC998 | CCMP1205 | RCC287 | RCC1871 | RCC2335 | RCC856 | RCC4434 | RCC3374 | RCC4656 | RCC696 | RCC999 | RCC4572 | RCC2339 |
|---|---|---|---|---|---|---|---|---|---|---|---|---|---|
| RCC998 | 0 | 6 | 6 | 6 | 6 | 5 | 30 | 16 | 18 | 18 | 18 | 18 | 18 |
| CCMP1205 | 6 | 0 | 0 | 0 | 0 | 1 | 32 | 13 | 16 | 16 | 16 | 16 | 16 |
| RCC287 | 6 | 0 | 0 | 0 | 0 | 1 | 32 | 13 | 16 | 16 | 16 | 16 | 16 |
| RCC1871 | 6 | 0 | 0 | 0 | 0 | 1 | 32 | 13 | 16 | 16 | 16 | 16 | 16 |
| RCC2335 | 6 | 0 | 0 | 0 | 0 | 1 | 32 | 13 | 16 | 16 | 16 | 16 | 16 |
| RCC856 | 5 | 1 | 1 | 1 | 1 | 0 | 32 | 12 | 15 | 15 | 15 | 15 | 15 |
| RCC4434 | 30 | 32 | 32 | 32 | 32 | 32 | 0 | 30 | 32 | 32 | 32 | 32 | 32 |
| RCC3374 | 16 | 13 | 13 | 13 | 13 | 12 | 30 | 0 | 3 | 3 | 3 | 3 | 3 |
| RCC4656 | 18 | 16 | 16 | 16 | 16 | 15 | 32 | 3 | 0 | 0 | 0 | 0 | 0 |
| RCC696 | 18 | 16 | 16 | 16 | 16 | 15 | 32 | 3 | 0 | 0 | 0 | 0 | 0 |
| RCC999 | 18 | 16 | 16 | 16 | 16 | 15 | 32 | 3 | 0 | 0 | 0 | 0 | 0 |
| RCC4572 | 18 | 16 | 16 | 16 | 16 | 15 | 32 | 3 | 0 | 0 | 0 | 0 | 0 |
| RCC2339 | 18 | 16 | 16 | 16 | 16 | 15 | 32 | 3 | 0 | 0 | 0 | 0 | 0 |

Figure S5

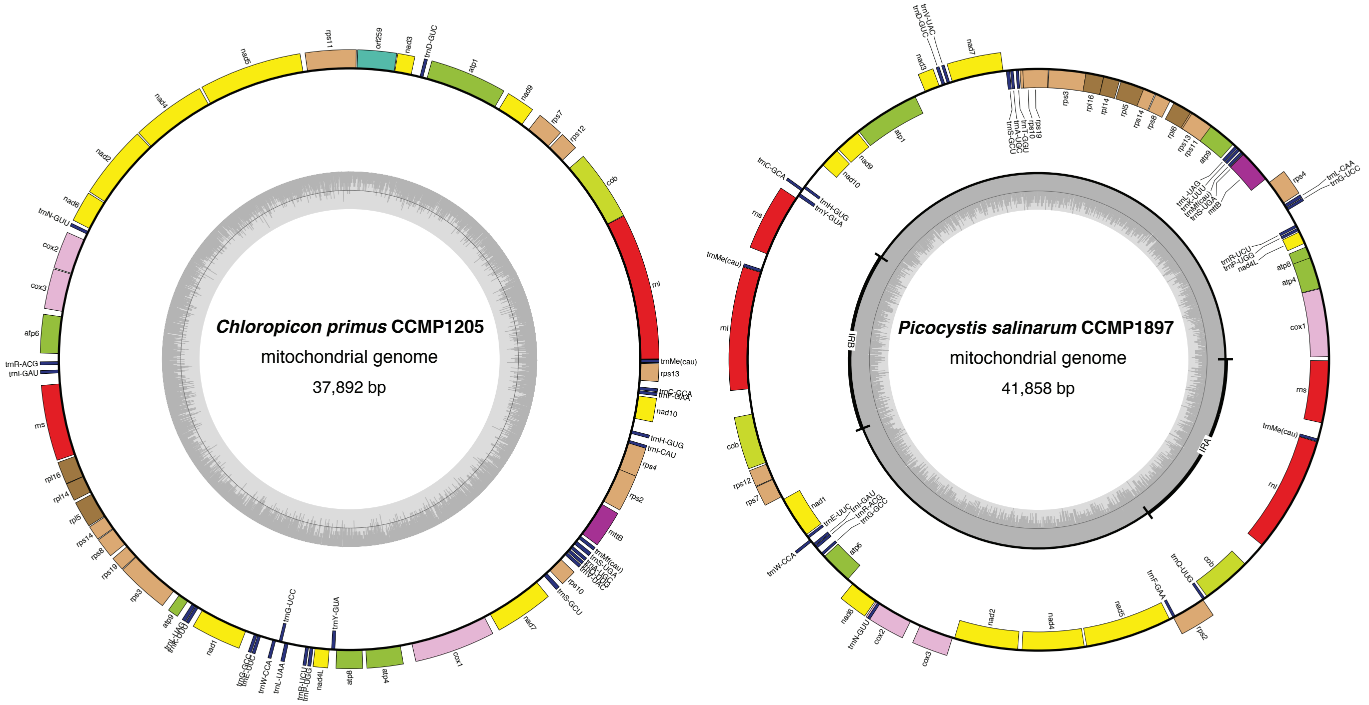


| Taxon \ Character | | rpl6 | I(cau) | L(caa) | L(uaa) | T(ggu) |
|---|---|---|---|---|---|---|
| 1 | Chloropicon_primus_CCMP1205_A2 | 0 | 1 | 0 | 1 | 0 |
| 2 | Picocystis_salinarum_CCMP1897_C | 1 | 0 | 1 | 0 | 1 |

Figure S6
