## Supplementary_tables_S1-S2 for "Tracing the evolution of the plastome and mitogenome in the Chloropicophyceae uncovered convergent tRNA gene losses and a variant plastid genetic code"

**Table S1**
Numbers of isoleucine (I) and leucine (L) codons in chloropicophycean and *Picocystis* plastomes and mitogenomes.

| | | Chloropicon | | | | | | | | Chloroparvula | | | | | Picocystis |
|---|---|---|---|---|---|---|---|---|---|---|---|---|---|---|---|
| Codon | AA | 998 (A1) | 1205 (A2) | 287(A3) | 1871 (A4) | 2335 (A4) | 856 (A5) | 4434 (A6) | 3374 (A7) | 4656 (B1) | 696 (B2) | 999 (B2) | 4572 (B3) | 4573 (B3) | 1897 (C) |
| **Plastomes** | | | | | | | | | | | | | | | |
| ATA | I | 111 | 90 | 103 | 71 | 73 | 95 | 147 | 109 | 254 | 25 | 30 | 25 | 47 | 189 |
| ATC | I | 220 | 199 | 237 | 221 | 225 | 251 | 175 | 192 | 98 | 150 | 152 | 150 | 106 | 150 |
| ATT | I | 751 | 818 | 758 | 793 | 788 | 741 | 834 | 797 | 1123 | 957 | 946 | 957 | 947 | 804 |
| CTA | L | 194 | 187 | 184 | 181 | 183 | 148 | 208 | 179 | 602 | 676 | 686 | 674 | 678 | 210 |
| CTC | L | 16 | 14 | 23 | 10 | 10 | 18 | 35 | 29 | 33 | 41 | 36 | 39 | 39 | 33 |
| CTG | L | 31 | 29 | 26 | 30 | 30 | 25 | 36 | 32 | 57 | 71 | 63 | 72 | 81 | 38 |
| CTT | L | 322 | 317 | 306 | 309 | 307 | 309 | 261 | 356 | 503 | 477 | 485 | 476 | 507 | 305 |
| TTA | L | 1167 | 1206 | 1178 | 1211 | 1206 | 1239 | 1156 | 1125 | 2 | 5 | 5 | 7 | 1 | 779 |
| TTG | L | 92 | 77 | 93 | 97 | 96 | 74 | 163 | 105 | 650 | 669 | 668 | 671 | 661 | 198 |
| **Mitogenomes** | | | | | | | | | | | | | | | |
| ATA | I | 160 | 93 | 92 | 65 | 66 | 111 | 69 | 33 | 28 | 3 | 3 | 4 | 1 | 2 |
| ATC | I | 114 | 156 | 172 | 195 | 193 | 134 | 144 | 116 | 111 | 47 | 46 | 48 | 65 | 150 |
| ATT | I | 312 | 317 | 297 | 305 | 307 | 312 | 342 | 402 | 490 | 439 | 445 | 439 | 411 | 32 |
| CTA | L | 162 | 205 | 219 | 213 | 220 | 168 | 166 | 116 | 363 | 369 | 366 | 365 | 292 | 15 |
| CTC | L | 39 | 50 | 40 | 39 | 31 | 40 | 45 | 36 | 20 | 13 | 12 | 13 | 17 | 328 |
| CTG | L | 25 | 27 | 37 | 37 | 37 | 39 | 54 | 24 | 17 | 39 | 38 | 41 | 55 | 679 |
| CTT | L | 417 | 459 | 438 | 413 | 412 | 400 | 283 | 317 | 263 | 179 | 181 | 180 | 260 | 64 |
| TTA | L | 409 | 375 | 355 | 376 | 374 | 405 | 370 | 412 | 2 | 2 | 3 | 2 | 1 | 2 |
| TTG | L | 71 | 25 | 35 | 36 | 36 | 63 | 144 | 104 | 387 | 424 | 426 | 425 | 459 | 227 |

**Table S2**
Number of SNPs and indels in compared plastomes and mitogenomes of *Chloropicon roscoffensis* (A4) and *Chloroparvula* sp. (B2 and B3) taxa.

| Compared taxa | RCC1871 (A4)<br>RCC2335 (A4) | RCC696 (B2)<br>RCC4572 (B3) | RCC696 (B2)<br>RCC999 (B2) | RCC999 (B2)<br>RCC4572 (B3) |
|---|---|---|---|---|
| **Plastomes** | | | | |
| Aligned nucleotides | 64533 | 70740 | 70732 | 70731 |
| Nucleotide identities | 64280 | 70530 | 69340 | 69328 |
| Indels (no.) | 5 | 10 | 25 | 25 |
| Nucleotides in indels (no.) | 17 | 14 | 92 | 90 |
| Nucleotide substitutions (no.) | 253 | 210 | 1392 | 1403 |
| Nucleotide substitution per site | 0.0039 | 0.0030 | 0.0197 | 0.0198 |
| **Mitogenomes** | | | | |
| Aligned nucleotides | 38147 | 35578 | 35409 | 35382 |
| nucleotide identities | 37707 | 35311 | 34610 | 34685 |
| Indels (no.) | 10 | 10 | 22 | 21 |
| Nucleotides in indels (no.) | 23 | 20 | 76 | 73 |
| Nucleotide substitutions (no.) | 440 | 267 | 799 | 697 |
| Nucleotide substitution per site | 0.0115 | 0.0075 | 0.0226 | 0.0197 |
